# Cytoplasmic and nuclear Sw-5b NLR act both independently and synergistically to dictate full host defense against tospovirus infection

**DOI:** 10.1101/2020.12.24.424293

**Authors:** Hongyu Chen, Xin Qian, Xiaojiao Chen, Tongqing Yang, Mingfeng Feng, Jing Chen, Ruixiang Cheng, Hao Hong, Ying Zheng, Yuzhen Mei, Danyu Shen, Yi Xu, Min Zhu, Xin Shun Ding, Xiaorong Tao

## Abstract

- Plant intracellular nucleotide binding-leucine-rich repeat (NLR) receptors play critical roles in mediating host immunity to pathogen attack. We use tomato Sw-5b::tospovirus as a model system to study the specific role of the compartmentalized plant NLR in dictating host defense against virus at different infection steps.
- We demonstrated here that tomato NLR Sw-5b translocates to cytoplasm and nucleus, respectively, to play different roles in inducing host resistances against *Tomato spotted wilt tospovirus* (TSWV) infection. The cytoplasmic Sw-5b functions to induce a strong cell death response to inhibit TSWV replication. This host response is, however, insufficient to block viral intercellular and long-distance movement. The nucleus-localized Sw-5b triggers a host defense that weakly inhibits viral replication but strongly impedes virus intercellular and systemic movement. Furthermore, the cytoplasmic and nuclear Sw-5b act synergistically to dictate full host defense to TSWV infection.
- We further demonstrated that the extended N-terminal *Solanaceae* domain (SD) of Sw-5b plays critical roles in cytoplasm/nucleus partitioning. Sw-5b nucleotide-binding leucine-rich repeat (NB-LRR) controls its cytoplasm localization. Strikingly, the SD but not coil-coil (CC) domain is crucial for Sw-5b receptor to translocate from cytoplasm to nucleus to trigger the immunity. The SD was found to interact with importins. Silencing both importin α and β expression disrupted Sw-5b nucleus translocation and host immunity against TSWV systemic infection.
- Collectively, our findings suggest that Sw-5b bifurcates disease resistances by cytoplasm/nucleus partitioning to block different infection steps of TSWV. The findings also identified a new regulatory role of extra domain of a plant NLR in mediating host innate immunity.

## Introduction

Plant innate immunity plays critical roles in host defense against pathogen invasions, and is triggered by cell-surface receptors or intracellular nucleotide-binding leucine-rich repeat (NLR) receptors (Soosaar *et al*., 2005; Dodds & Rathjen, 2010; Cui *et al*., 2015; Li *et al*., 2015; Jones *et al*., 2016; Kourelis & van der Hoorn, 2018; Kapos *et al*., 2019; van Wersch, 2020). Plant intracellular NLRs are the largest classes of resistance proteins that function to detect pathogen effectors, and to activate host immunity upon pathogen attack (Caplan, J *et al*., 2008; Takken & Goverse, 2012; Li *et al*., 2015; Jones *et al*., 2016; Kourelis & van der Hoorn, 2018; Kapos *et al*., 2019). Plant NLRs typically contain an N-terminal domain, a central nucleotide-binding domain (NB), a nucleotide-binding adaptor (ARC domain shared by Apaf-1, certain resistance proteins, and CED-4), and a C-terminal leucine-rich repeat (LRR) domain (Ea & Jones, 1998; Jones *et al*., 2016; Ma *et al*., 2018; Wang *et al*., 2019a; Wang *et al*., 2019b; Ma *et al*., 2020). Based on the differences among the N-terminal domains, plant NLRs can be further divided into two main categories, known as the coiled-coil NLR (refers to as CNL) category and the Toll/interleukin-1 receptor NLR (TNL) category (Meyers *et al*., 2003; Collier & Moffett, 2009; Qi & Innes, 2013). The CC- or the TIR-domain-bearing NLRs have distinct genetic requirements and can regulate different functions in the downstream of defense signaling (Collier & Moffett, 2009; Qi & Innes, 2013; Horsefield *et al*., 2019; Jubic *et al*., 2019; van Wersch & Li, 2019; Wan *et al*., 2019).

In addition to classical domains, non-canonical domains were frequently found to integrate into certain NLRs. The additional domain, called BED, was first found in 32 poplar NLR proteins (Germain & Seguin, 2011). This extra BED domain was also found in nine rice NLRs (Das *et al*., 2014). The RATX1/HMA domain in the rice NLRs RGA5 and Pik - 1 was found to act as integrated decoys to detect the cognate pathogen effectors (Kanzaki *et al*., 2012; Cesari *et al*., 2013; Cesari *et al*., 2014). The WRKY domain on Arabidopsis thaliana NLR RRS1 was further found to function as an integrated decoy that recognizes the effectors AvrRps4 and PopP2 (Le Roux *et al*., 2015; Sarris *et al*., 2015). Genome-wide analyses of plant NLR receptors revealed that about 3.5 % of the NLRs carried specific non-canonical domains (Cesari *et al*., 2014; Kroj *et al*., 2016; Sarris *et al*., 2016), and some of these non-canonical domains were shown to be targeted by pathogen effectors during pathogen infections (Sarris *et al*., 2016). However, molecular functions of most non-canonical domains in plant NLRs remain largely unexplored.

Translocations of plant NLRs into proper subcellular compartments are critical for the induction of innate immunity (Cui *et al*., 2015; van Wersch, 2020). Multiple plant NLRs and immune regulators, including tobacco N, Arabidopsis snc1, RRS1/RPS4, barley MLA10, and Arabidopsis EDS1, NPR1, have been shown to accumulate in both cytoplasm and nucleus, and for several nucleocytoplasmic NLRs accumulation in nucleus is required for triggering host resistance to pathogen infections (Deslandes *et al*., 2003; Burch-Smith *et al*., 2007; Shen *et al*., 2007; Wirthmueller *et al*., 2007; Tasset *et al*., 2010; Bai *et al*., 2012; Inoue *et al*., 2013; Padmanabhan *et al*., 2013). Wheat Sr33, a homolog of barley MLA10, however, was reported to accumulate in cytoplasm to induce host resistance against stem rust pathogen (Cesari *et al*., 2016). For potato Rx, a balanced cytoplasm and nucleus accumulation of Rx is needed to induce the host immunity (Slootweg *et al*., 2010; Tameling *et al*., 2010). Other studies have shown that *Arabidopsis* Rpm1 (Gao *et al*., 2011), RPS2 (Axtell & Staskawicz, 2003), RPS5 (Qi *et al*., 2012), rice Pit (Takemoto *et al*., 2012), and tomato Tm-2^2^ (Chen *et al*., 2017; Wang *et al*., 2020) need to associate with plasma membrane in order to trigger cell death and host immunity. Latest studies have shown that the activated *Arabidopsis* ZAR1 can bind to cellular membrane, leading to a membrane leakage followed by cell death and host immunity (Wang *et al*., 2019a; Wang *et al*., 2019b). Flax L6 and M have been shown to accumulate in both Golgi apparatus and tonoplast, and these compartmentalized localizations are necessary for the induction of host resistance (Kawano *et al*., 2014). The re-distribution of potato R3a from cytosol to endosomal compartments is crucial for the induction of host resistance to *Phytophthora infestans* infection (Engelhardt *et al*., 2012). Different plant NLRs have diverse subcellular localizations for their proper functions. However, how the compartmentalized plant NLRs specifically dictate defense signaling remains largely unknown.

Tomato spotted wilt tospovirus (TSWV) is one of most destructive plant NSVs, infecting more than 1000 plant species, and causes crop losses more than one billion US dollars annually worldwide (Kormelink *et al*., 2011; Scholthof *et al*., 2011; Oliver & Whitfield, 2016). Tomato NLR Sw-5b confers strong resistance to TSWV infection and has been widely used in tomato breeding projects to produce tospovirus resistant tomato cultivars (Brommonschenkel *et al*., 2000; Spassova M I, 2001; Turina *et al*., 2016; Zhu *et al*., 2019). Upon recognition of TSWV movement protein, NSm, Sw-5b can trigger a hypersensitive response (HR), which typically associated with localized cell death (Lopez *et al*., 2011; Hallwass *et al*., 2014; Peiro *et al*., 2014; De Oliveira *et al*., 2016; Zhao *et al*., 2016; Leastro *et al*., 2017). Tospoviruses are divided into American and Euro-Asia type based on their geographic distribution and amino acid sequence identity of viral nucleocapsid protein. We have previously shown that Sw-5b can confer a broad-spectrum resistance to American type tospoviruses, including TSWV, through recognition of a conserved 21 amino acid PAMP-like region in the viral movement protein NSm (Zhu *et al*., 2017). Sw-5b carries an extended N-terminal *Solanaceae* domain (SD), a CC domain, a NB-ARC domain, and a LRR domain (Brommonschenkel *et al*., 2000; Spassova M I, 2001; Lukasik-Shreepaathy *et al*., 2012). Similar SDs have also been found in the Mi-1.2, R8, Rpi-blb2, and Hero (Milligan *et al*., 1998; Vos *et al*., 1998; Ernst *et al*., 2002; van der Vossen *et al*., 2005; Lukasik-Shreepaathy *et al*., 2012; Vossen *et al*., 2016). More recently, Seong and others reported that the extended CNL has been evolved initially in the ancestor of *Asterids* and *Amaranthaceae*, predated the *Solanaceae* family (Seong *et al*., 2020). In the presence of the extended N-terminal SD, Sw-5b is in an autoinhibited state through multilayered interactions between SD, CC, NB-ARC, and LRR domains (Chen *et al*., 2016). For activation, the extra SD also recognizes NSm. Sw-5b adopts a two-step NSm recognition strategy through SD and then LRR domain (Li *et al*., 2019). This two-step recognition mechanism significantly enhances the sensitivity of the detection on TSWV NSm (Li *et al*., 2019). Although Sw-5b is known to localize in both cytoplasm and nucleus (De Oliveira *et al*., 2016), the biological roles of the cytoplasm- and the nucleus-accumulated Sw-5b in host immunity signaling are unknown.

In this study, we investigated the subcellular distribution pattern of Sw-5b and the functions of the compartmentalized Sw-5b in the induction of host immunity to TSWV infection. We determined here that cytoplasm- and nucleus-accumulated Sw-5b functions differently in inducing host defense response to inhibit multiple tospovirus infection steps. The cytoplasmic Sw-5b can induce a strong cell death response to suppress TSWV replication, whereas the nucleus-accumulated Sw-5b can induce a strong defense against viral intercellular movement and systemic infection. The combination of cytoplasmic and nuclear Sw-5b induces a synergistic and strong plant immunity against tospovirus infection. We also found that the extended SD functions as the key regulator for this critical intracellular translocation. The SD was also found to interact with importins α and β to mediate Sw-5b nucleus translocation, and to confer the full host immunity against tospovirus infection.

## Materials and Methods

### Plasmid construction

p2300S-YFP-Sw-5b was from a previously described source (Chen *et al*., 2016). Different domains of Sw-5b were PCR-amplified from p2300S-Sw-5b (Chen *et al*., 2016) and cloned individually behind the *YFP* gene in the p2300S vector using a two-step overlap PCR procedure as described (Li *et al*., 2019). All the primers used in this study are listed in Table S1.

To visualize the subcellular localization patterns of various fusion proteins, a SV40 T-Ag-derived nuclear localization signal (NLS, QPKKKRKVGG) (Lanford & Butel, 1984) or a PK1 nuclear export signal (NES, NELALKLAGLDINK) (Wen *et al*., 1995) was fused to the N-terminus of YFP-Sw-5b or the C-terminus of NSm-YFP, as described (Kong *et al*., 2017), to produce pNES-YFP-Sw-5b, pNLS-YFP-Sw-5b, pNSm-YFP-NES, and pNSm-YFP-NLS, respectively. In addition, YFP-Sw-5b and NSm-YFP were fused individually with a mutant NLS (nls, QPKKTRKVGG) or a mutant NES (nes, NELALKAAGADANK) to produce pnes-YFP-Sw-5b, pnls-YFP-Sw-5b, pNSm-YFP-nes, and pNSm-YFP-nls. The constructs were then transformed individually into *Agrobacterium tumefaciens* strain GV3101 cells.

### Transient gene expression, stable plant transformation, and virus inoculation

*Nicotiana benthamiana* were grown in soil in pots inside a greenhouse maintained at 25°C and a 16 h light/8 h dark photoperiod. Six-to-eight week-old *N. benthamiana* plants were used for various assays. Transient gene expression assays were performed in *N. benthamiana* leaves through agro-infiltration using Agrobacterium cultures carrying specific expressing constructs as described previously (Feng *et al*., 2016; Ma *et al*., 2017). Transgenic *N. benthamiana* lines expressing YFP-Sw-5b or its derivatives were made using constructs with a 35S promoter or a Sw-5b native promoter via a standard leaf-disc transformation method (Chen *et al*., 2016). The resulting transgenic *N. benthamiana* lines were named as NES-YFP-Sw-5b, NLS-YFP-Sw-5b, nes-YFP-Sw-5b, nls-YFP-Sw-5b, YFP-Sw-5b, and EV (transformed with an empty vector), respectively. Inoculation of transgenic *N. benthamiana* plants with TSWV was done by rubbing plant leaves with TSWV-YN isolate-infected crude saps as described (Zhu *et al*., 2017). TRV-mediated VIGS in *N. benthamiana* plants was done as described (Ma *et al*., 2015). The agro-infiltrated or the virus-inoculated plants were growning inside a growth chamber maintained at 25/23 °C (day/night) with a 16/8 h light and dark photoperiod.

### Particle bombardment

The particle bombardment is described (Feng et al., 2016). Briefly, 60 mg Tungsten M-10 microcarrier (Bio-RAD) was placed into a 1.5 ml Eppendorf tube with 1 mL 70% ethanol. The tube was vortexes for 3 minutes, and then stood at room temperature for 15 minutes. After centrifuge at low speed for 5 seconds, the supernatant was removed and the pellet was rinsed with 70% ethanol for 3 times. One mL 50% sterile glycerol solution was added and divided Tungsten M-10 microcarrier into 50 µl. Five µg pRTL2-YFP, pRTL2-YFP-Sw-5b or pRTL2-YFP-Sw-5bD857V plasmid DNA, 50 µl of 2.5 M CaCl2, and 20 µl of 0.1 M spermidine, respectively were added and mixed with microcarrier. After centrifuge at low speed for 5 seconds and the supernatant removed. The pellet was resuspended in 200 µl 70% ethanol and centrifugation as described above. Use 48 µl of 100% ethanol to resuspend the tungsten particle::plasmid DNA complexes, and load 15 48 µl mixture onto the center of carrier (Bio-RAD), air dry, and use He/1000 particle transport system (BIO-RAD) to bombard tomato leaves harvested from 3- or 4-week-old of Money Marker. The bombarded leaves were incubated in Petri dishes for 24 hours at 25°C followed with Confocal Microscope analysis.

### Trypan blue staining

*N. benthamiana* leaves were harvested at 3 days post agro-infiltration (dpai) and boiled for 5 min in a 1.15:1 (v/v) mixed ethanol and trypan blue staining solution (10 g phenol, 10 mL glycerol, 10 mL lactic acid, and 20 mg trypan blue in 10 mL distilled water). The stained leaves were then de-stained in a chloral hydrate solution (2.5 g per mL distilled water) as described (Bai *et al*., 2012).

### Electrolyte leakage assay

Electrolyte leakage assay was performed as previously described (Mittler *et al*., 1999; Zhu *et al*., 2017) with slight modifications. Briefly, five leaf discs (9 mm in diameter each) were taken from the agro-infiltrated leaves per treatment and at various dpai. The harvested leaf discs from a specific treatment were floated on a 10 mL distilled water for 3 h at room temperature (RT), and the conductivity of each bathing water was measured (referred to as value A) using a Multiparameter Meter as instructed (Mettler Toledo, Zurich, Switzerland). After the first measurement, the leaf discs were returned to the bathing water and incubated at 95°C for 25 min. After cooling down to RT, the conductivity of each bathing sample was measured again (referred to as value B). The ion leakage was expressed as the ratio determined by value A/value B × 100. The mean value and standard error of each treatment were calculated using the data from three biological replicates per treatment at each sampling time point.

### Confocal laser scanning microscopy

Tissue samples were collected from the leaves of transiently expressing YFP-Sw-5b or one of the fusion proteins at 24–36 hours post agro-infiltration (hpai). The collected tissue samples were mounted in water between a glass slide and a coverslip. Images of individual samples were captured under a Carl Zeiss LSM 710 confocal laser scanning microscope. YFP fusions were excited at 488 nm and the emission was captured at 497–520 nm. The resulting images were further processed using the Zeiss 710 CLSM software followed by the Adobe Photoshop software (San Jose, CA, USA).

### Nucleus and cytoplasm fractionations

*N. benthamiana* leaf tissues (1 g per sample), representing a specific treatment, were collected at 24 hpai, frozen in liquid nitrogen, ground to fine powders, and then homogenized in 2 mL (per sample) extraction buffer 1 (20 mM Tris-HCl, pH 7.5, 20 mM KCl, 2.5 mM MgCl2, 2 mM EDTA, 25% glycerol, 250 mM sucrose, 1×Protease Inhibitor Cocktail, and 5 mM DTT). The resulting lysate was filtered through 30 μm filters to remove debris, and the filtrate was centrifuged at 2,000 × *g* for 5 minutes to pellet nuclei. The supernatant from a sample was transferred into a new tube and centrifuged at 10,000 × *g* for 10 min. The resulting supernatant was used as the cytoplasm fraction. The nuclei containing pellet was resuspended in 5 mL extraction buffer 2 (20 mM Tris-HCl, pH 7.4, 25% glycerol, 2.5 mM MgCl2, and 0.2% Triton X-100), centrifuged for 10 min at 2,000 × *g* followed by 4-6 cycles of resuspension and centrifugation as described above. The resulting pellet was resuspended again in 500 μl extraction buffer 3 (20 mM Tris-HCl, pH 7.5, 0.25 M sucrose, 10 mM MgCl2, 0.5% Triton X-100, and 5 mM β-mercaptoethanol). The nuclei fraction was carefully layered on the top of 500 mL extraction buffer 4 (20 mM Tris-HCl, pH 7.5, 1.7 M sucrose, 10 mM MgCl2, 0.5% Triton X-100, 1×Protease Inhibitor Cocktail, and 5 mM β-mercaptoethanol), and then centrifuged at 16,000 × *g* for 1 h. The resulting pellet was resuspended in 500 μL extraction buffer 1 and stored at −80°C until use or used immediately for SDS-PAGE assays. All the processes were performed on ice or at 4°C. In this study, actin and histone H3 were used as the cytoplasmic and the nuclear markers, respectively.

### Western blot, co-immunoprecipitation and mass spectrometry analysis

Western blot and co-immunoprecipitation assays were performed as described (Zhu *et al*., 2017). Briefly, agro-infiltrated leaf samples (1 g per sample) were harvested and homogenized individually in pre-chilled mortars with pestles in 2 mL extraction buffer [10% (v/v) glycerol, 25 mM Tris, pH 7.5, 1 mM EDTA, 150 mM NaCl, 10 mM DTT, 2% (w/v) polyvinylpolypyrrolidone, and 1 × protease inhibitor cocktail (Sigma, Shanghai, China)]. Each crude slurry was transferred into a 2 mL Eppendorf tube, and spun for 2 min at full speed in a refrigerated microcentrifuge. The supernatant was transferred into a clean 1.5 mL Eppendorf tube and spun for 10 min at 4°C. For Western blot assays, 50 μL supernatant from a sample was mixed with 150 μL Laemmli buffer, boiled for 5 min, and analyzed in SDS-PAGE gels through electrophoresis. For immunoprecipitation assays, 1 mL supernatant was mixed with 25 μL GFP-trap agarose beads (ChromoTek, Planegg-Martinsried, Germany), incubated for 2 h at 4°C on an orbital shaker, and then pelleted through low speed centrifugation. The blots were probed with a 1:2,500 (v/v) diluted anti-YFP antibody or other specific antibodies followed a 1:10,000 (v/v) diluted horseradish peroxidase (HRP)-conjugated goat anti-rabbit or a goat anti-mouse antibody (Sigma-Aldrich, St. Louis, MO, USA). The detection signal was developed using the ECL substrate kit as instructed (Thermo Scientific, Hudson, NH, USA).

For mass spectrometry analysis, the immunoprecipitation samples of YFP-SD and SD without tag were processed by The Beijing Genomics Institute (BGI) for mass spectrometry analysis. The immunoprecipitation samples of YFP-Sw-5b and Sw-5b without tag were processed by Applied Protein Technology in Shanghai. Database searches were performed using the Mascot search engine against *N. benthamiana*. (https://solgenomics.net/organism/Nicotiana_benthamiana/genome).

### RT-PCR detection of TSWV infection

Total RNA was extracted from TSWV-inoculated *N. benthamiana* plant leaves using an RNA Purification Kit (Tiangen Biotech, Beijing, China), and then treated with RNase-free DNase I (TaKaRa, Dalian, China). First-strand cDNA was synthesized using a TSWV-specific primer (S3 Table). PCR reactions were as follows: initial denaturation at 94°C for 2 min followed by 35 cycles of 94°C for 30 s, 52°C for 30 s, and 72°C for 1 min. The final extension was 72°C for 10 min. The resulting PCR products were visualized in 1.0% (w/v) agarose gels through electrophoresis.

## Results

### Determination of Sw-5b subcellular localization pattern

Expression of YFP-Sw-5b in *N. benthamiana* leaves resulted in a strong HR cell death as well as Sw-5b (Chen *et al*., 2016; Zhu *et al*., 2017). To investigate the subcellular localization pattern of Sw-5b, we transiently expressed YFP and YFP-Sw-5b in *N. benthamiana* leaves, respectively, through agro-infiltration. Confocal Microscopy results showed that the YFP-Sw-5b fusion accumulated in both cytoplasm and nucleus in *N. benthamiana* leaf cells (Fig. 1b, middle image). This subcellular localization pattern was similar to that of YFP (Fig. 1b, left image). When a D857V mutation, which keeps Sw-5b in an autoactivated state (Chen *et al*., 2016), was introduced into Sw-5b to produce pYFP-Sw-5b^D857V^ and expressed in *N. benthamiana* leaves, the mutant YFP-Sw-5b^D857V^ fusion also accumulated in both cell cytoplasm and nucleus (Fig. 1b, right image). We also making a construct pNativePro::YFP-Sw-5b expressing YFP-Sw-5b under native Sw-5b promoter. However, the expression of YFP-Sw-5b by native Sw-5b promoter is too low to detect green fluorescence signal.

**Fig. 1.**
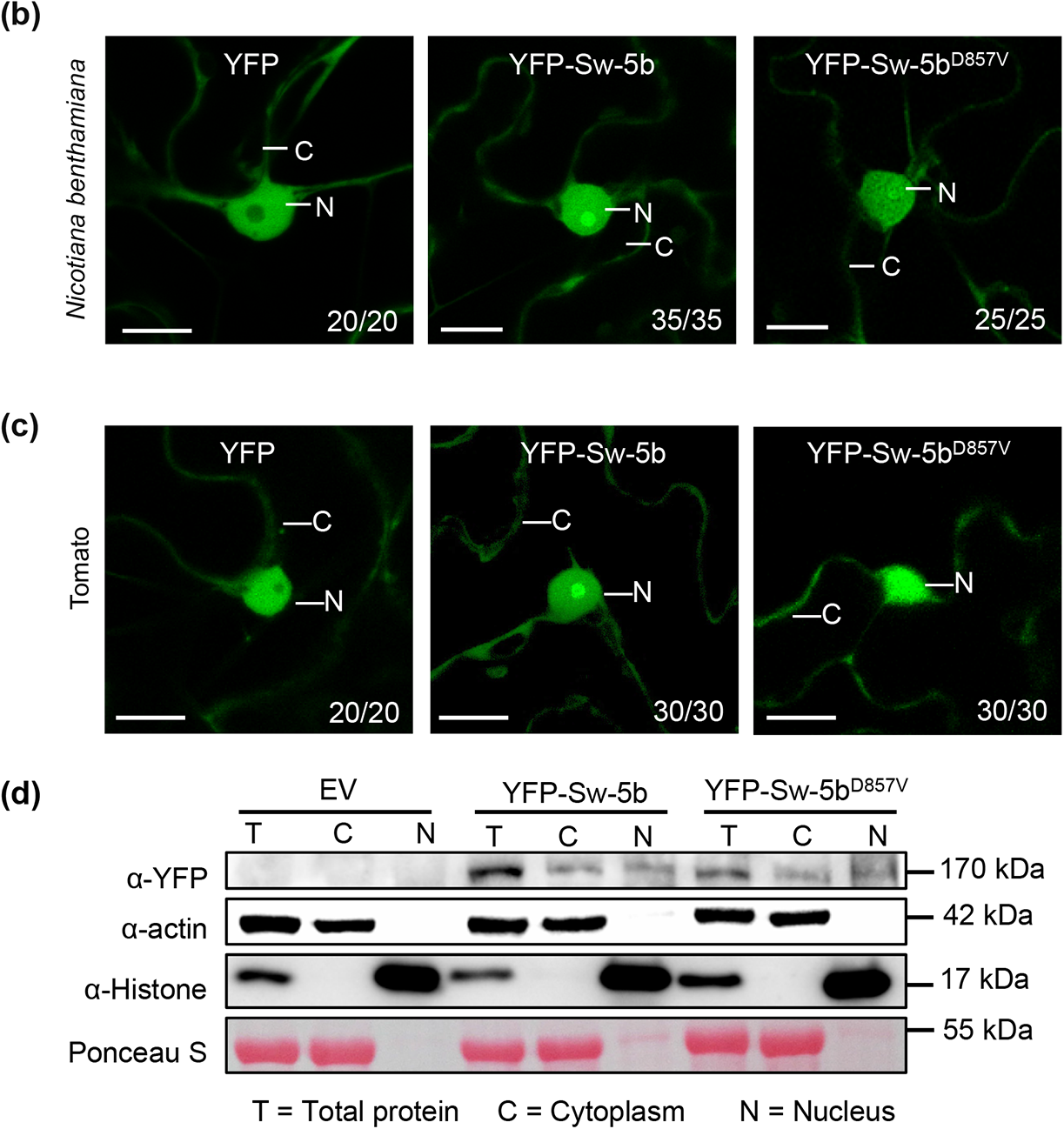
Subcellular localization of Sw-5b in *Nicotiana benthamiana* and tomato leaf cells. (a) Schematic diagrams of Sw-5b. (b) Subcellular localizations of free YFP (left), YFP-Sw-5b (middle) and autoactive YFP-Sw-5b^D857V^ mutant (right) in *N. benthamiana* leaf cells at 24 hours post agro-infiltration (hpi). (c) Subcellular localization of free YFP (left), YFP-Sw-5b (middle) and autoactive YFP-Sw-5b^D857V^ mutant (right) in tomato leaf cells at 24 hpi. N nucleus, and C cytoplasm inside the cell are indicated. Bar = 10 μm. (d) Nucleocytoplasmic partitioning analysis of YFP-Sw-5b and autoactive YFP-Sw-5b^D857V^. Total lysate (T) from p2300S empty vector (EV), YFP-Sw-5b or YFP-Sw-5b^D857V^ expressing leaves were fractionated into cytoplasm and nucleus, and analyzed by immunoblots using antibodies against YFP. The actin and histone were used as a cytoplasm marker and nucleus marker, respectively, in the fractionation analysis. Ponceau S staining was also used as cytoplasm marker.

To investigate the subcellular localization pattern of Sw-5b in tomato leaf cells, we transiently expressed YFP, YFP-Sw-5b, and YFP-Sw-5b^D857V^, respectively, through particle bombardment. Confocal Microscopy results showed that these three proteins exhibited the same subcellular localization pattern as that in the *N. benthamiana* leaf cells (Fig. 1c).

To further confirm the above results, we harvested *N. benthamiana* leaves expressing YFP-Sw-5b or YFP-Sw-5b^D857V^ and analyzed by cytoplasm and nucleus fractionation assay. Leaf samples agro-infiltrated with the empty vector (p2300S) were also harvested and used as controls. Analyses of total protein, cytoplasm fractions, and nucleus fractions from these harvested leaves using Western blot assays showed that both YFP-Sw-5b and YFP-Sw-5b^D857V^ were accumulated in the cytoplasm and nucleus (Fig. 1d).

### Sw-5b recognizes TSWV NSm in the cytoplasm

TSWV NSm is known to reside in cytoplasm and plasmodesmata, but not in nucleus (Kormelink *et al*., 1994; Feng *et al*., 2016). To determine where Sw-5b can recognize TSWV NSm, we fused a NES, a nes, a NLS or a nls signal peptide to the C-terminus of NSm-YFP to produce NSm-YFP-NES, NSm-YFP-nes, NSm-YFP-NLS, and NSm-YFP-nls, respectively. Transient expressions of these fusions in *N. benthamiana* leaves showed that NSm-YFP-NES accumulated exclusively in the cytoplasm, while NSm-YFP-NLS accumulated in the nucleus (Fig. S1a). As expected, NSm-YFP-nes and NSm-YFP-nls showed the same accumulation pattern as that of NSm-YFP (Fig. S1a). When Sw-5b was co-expressed with one of the above four fusions in *N. benthamiana* leaves through agro-infiltration, the leaf tissues co-expressing Sw-5b and NSm-YFP-NES (Sw-5b + NSm-YFP-NES), Sw-5b and NSm-YFP-nes (Sw-5b + NSm-YFP-nes), or Sw-5b and NSm-YFP-nls (Sw-5b + NSm-YFP-nls) developed a strong HR cell death (Fig. S1b). In contrast, the leaf tissues co-expressing Sw-5b and NSm-YFP-NLS (Sw-5b + NSm-YFP-NLS) did not. Western blot assays using a YFP specific antibody confirmed that all the assayed proteins were expressed in the infiltrated tissues (Fig. S1c), indicating that Sw-5b recognizes NSm in the cytoplasm.

### Sw-5b activity in cell death induction is enhanced in the cytoplasm but suppressed in the nucleus

To investigate the roles of the cytoplasmic and nuclear Sw-5b in the induction of cell death and host immunity, we produced constructs to express YFP-Sw-5b, NLS-YFP-Sw-5b, nls-YFP-Sw-5b, NES-YFP-Sw-5b, and nes-YFP-Sw-5b, respectively, and then tested their abilities to elicit cell death and host immunity to tospovirus infection. Transient expressions of these fusions in *N. benthamiana* leaves showed that NES-YFP-Sw-5b accumulated exclusively in the cytoplasm, while NLS-YFP-Sw-5b accumulated only in the nucleus (Fig. 2a). In addition, nes-YFP-Sw-5b and nls-YFP-Sw-5b showed the same accumulation pattern as that shown by YFP-Sw-5b. We then tested cell death induction through co-expressions of NSm and YFP-Sw-5b (NSm + YFP-Sw-5b), NSm and NES-YFP-Sw-5b (NSm + NES-YFP-Sw-5b), NSm and nes-YFP-Sw-5b (NSm + nes-YFP-Sw-5b), NSm and NLS-YFP-Sw-5b (NSm + NLS-YFP-Sw-5b), or NSm and nls-YFP-Sw-5b (NSm + nls-YFP-Sw-5b) in *N. benthamiana* leaves through agro-infiltration. Results of this study showed that the NSm + NES-YFP-Sw-5b-induced cell death was stronger than that induced by NSm + nes-YFP-Sw-5b or NSm + nls-YFP-Sw-5b co-expression (Fig. 2b). In addition, the cell death induced by NSm + NLS-YFP-Sw-5b co-expression was suppressed (Fig. 2b). Western blot results showed that the stronger cell death caused by NSm + NES-YFP-Sw-5b co-expression was not due to a greater accumulation of NES-YFP-Sw-5b in the leaves (Fig. 2c). The ion leakage assay results (Fig. 2d) agreed with the phenotype observation results, and indicated that co-expression of NSm + NES-YFP-Sw-5b in leaves lead to a greater ion leakage compared with that induced by the co-expression of NSm + nes-YFP-Sw-5b at 24 and 48 hours post agro-infiltration (hpai). The ion leakage caused by the co-expression of NSm + NLS-YFP-Sw-5b was significantly weaker than that caused by the co-expression of NSm + nls-YFP-Sw-5b (Fig. 2d).

**Fig. 2.**
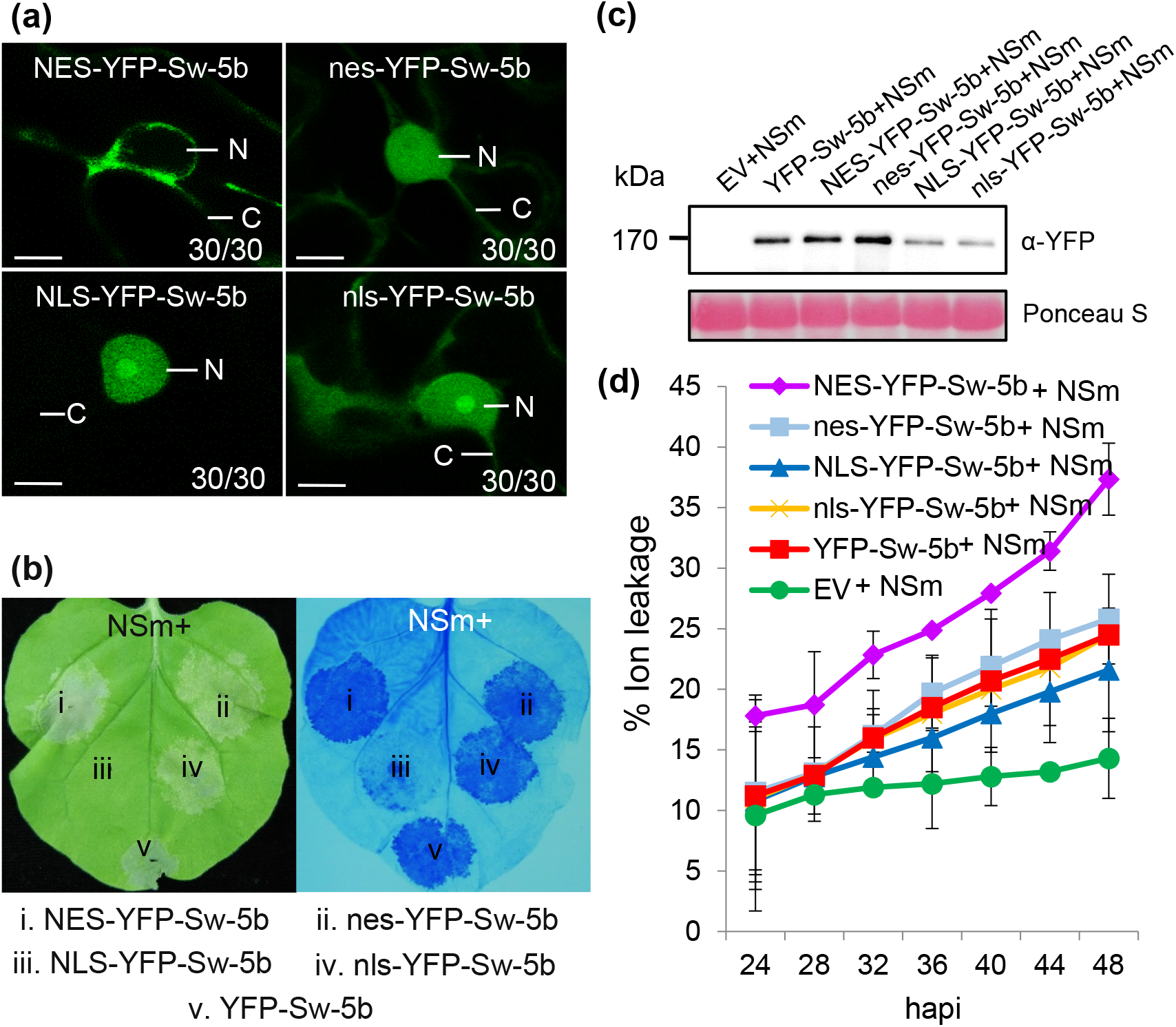
Effect of Sw-5b subcellular localization pattern on HR induction. (a) Confocal images of *N. benthamiana* leaf cells transiently expressing NES-YFP-Sw-5b, nes-YFP-Sw-5b, NLS-YFP-Sw-5b or nls-YFP-Sw-5b fusion. The images were taken at 24–36 hpi. N nucleus and C cytoplasm (c). Bar = 10 μm. (b) Induction of HR in *N. benthamiana* leaf tissues co-expressing NSm and one of the five Sw-5b fusion proteins. The infiltrated *N. benthamiana* leaf was photographed at 3 dpi (left image). Induction of HR in the infiltrated tissues were visualized using a trypan blue staining method (right image). (c) Immunoblot analysis of NES-YFP-Sw-5b, nes-YFP-Sw-5b, NLS-YFP-Sw-5b, and nls-YFP-Sw-5b expressions in the infiltrated *N. benthamiana* leaf tissues. These fusion proteins were enriched using the GFP-Trap beads prior to SDS-PAGE, and the blot was probed using an YFP specific antibody. Ponceau-S staining was used to estimate sample loadings. (d) Time course analysis of ion leakage in Nicotiana benthamiana leaves co-expressing NSm with one of the five Sw-5b fusion proteins. Measurements were performed at 4 h intervals starting from 24 to 48 hpi. Error bars (SEs) were calculated using the results from three biological replicates per treatment collected at each time point.

### Cytoplasmic Sw-5b induces a strong host defense against tospovirus replication

Virus infection in plant starts with virus replication in the initially infected cells followed by spreading into adjacent cells for further infection. To monitor tospovirus replication in plant cells, we recently developed a TSWV mini-replicon-based reverse genetic system (Feng et al., 2020). In this study, co-expression of TSWV mini-replicon SR(+)eGFP, L(+)opt (with a codon usage optimized RdRp), VSRs and NSm resulted in a cell-to-cell movement of SR(+)eGFP. In contrast, co-expression of SR(+)eGFP, L(+)opt, VSRs, and NSm^H93A&H94A^ mutant, a defective movement protein but can be recognized by Sw-5b to cause a strong HR (Li *et al*., 2009; Zhao *et al*., 2016), in cells resulted in the expression of SR(+)eGFP in only single cells (Fig. S2), thus dissecting the viral replication from viral cell-to-cell movement. We then co-expressed SR(+)eGFP, L(+)opt, VSRs, NSm^H93A&H94A^ mutant and one of the five proteins (i.e., Sw-5b, NES-Sw-5b, nes-Sw-5b, NLS-Sw-5b, nls-Sw-5b) in *N. benthamiana* leaves. Leaves co-expressing SR(+)eGFP, L(+)opt, VSRs, NSm^H93A&H94A^ mutant and p2300 (empty vector, EV) were used as controls. The results showed that in the presence of Sw-5b or one of its derivatives, the expression of SR(+)eGFP was strongly suppressed compared with that expressed in the presence of EV (Fig. 3a). It is noteworthy that the expression of SR(+)eGFP was less inhibited in the presence of NLS-Sw-5b (Fig. 3a). Western blot result indicated that the GFP accumulation of SR(+)eGFP was strongly inhibited in the presence of Sw-5b, NES-Sw-5b, nes-Sw-5b or nls-Sw-5b compared with that expressed in the presence of NLS-Sw-5b or EV (Fig. 3b). This finding indicates that the cytoplasmic Sw-5b can inhibit SR(+)eGFP expression, possibly through induction of a host defense against TSWV replication.

**Fig. 3.**
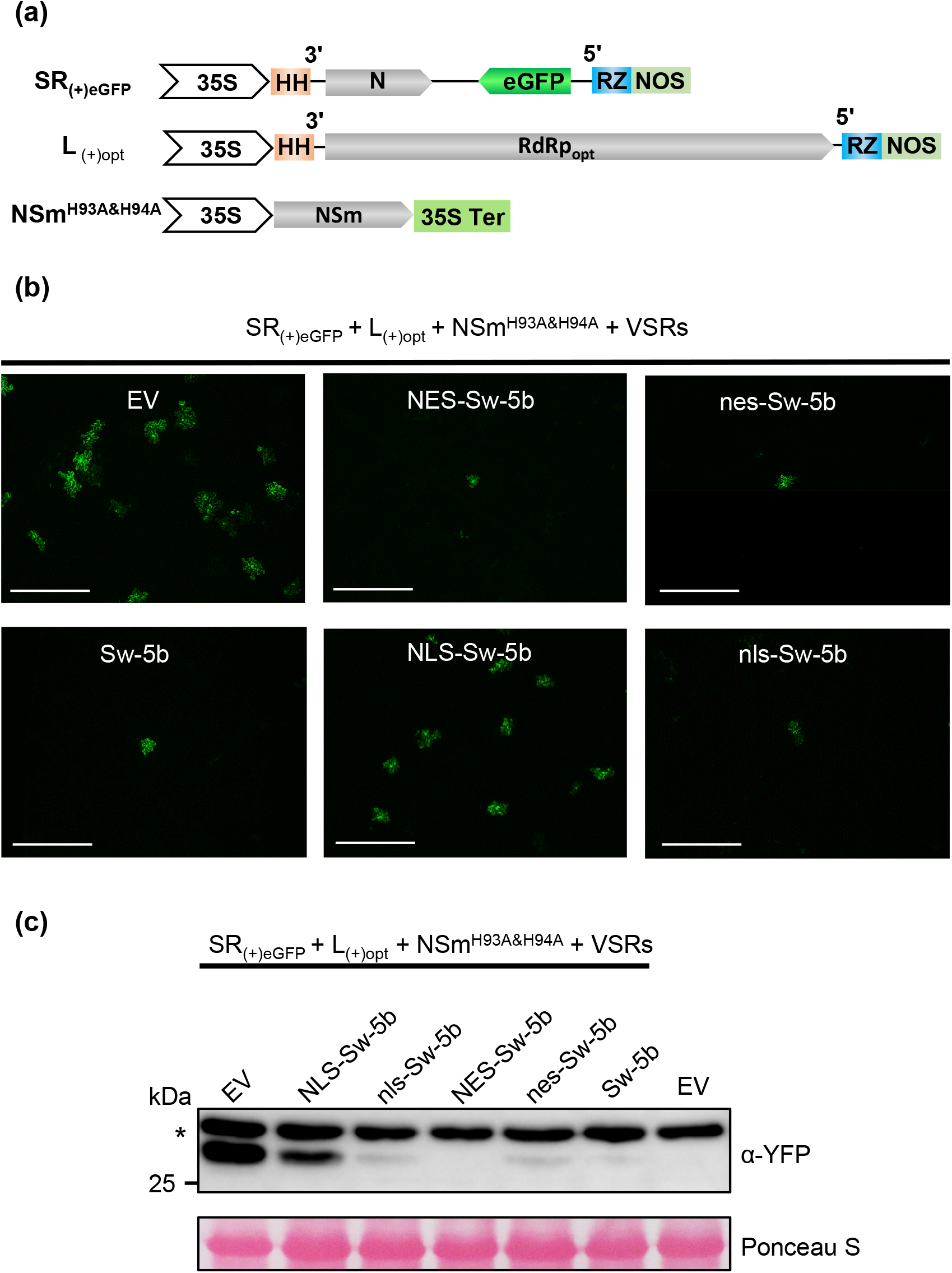
The effect of cytoplasm- and nucleus-targeted Sw-5b on viral replication. (a) Schematic representation of binary constructs to express TSWV SR(-)eGFP mini-genome replicon, TSWV L RNA segment containing an optimized RdRp and NSm^H93A&H94A^ mutant that defected in viral movement. Minus sign (-) and 5′ to 3′ designation represent the negative (genomic)-strand of tospovirus RNA. 35S: a double 35S promoter; HH: hammerhead ribozyme; RZ: hepatitis delta virus (HDV) ribozyme; NOS: nopaline synthase terminator; 35S Ter: a 35S transcription terminator. (b) Accumulation of eGFP fluorescence in *N. benthamiana* leaves co-expressing p2300S empty vector (EV), Sw-5b, NES-Sw-5b, nes-Sw-5b, NLS-Sw-5b, or nls-Sw-5b with TSWV SR(-)eGFP, L, and NSm^H93A&H94A^ at 4 days post infiltration (dpi) viewed with a fluorescence microscope. Bar represents 400 μm. (c) Immunoblot analysis of expression of eGFP proteins in leaves shown in panel (b) using specific antibodies against YFP. Ponceau S staining of rubisco large subunit is shown for protein loading control.

### Sw-5b induces a host defense against viral NSm intercellular movement

In our previous study, we used pmCherry-HDEL//NSm-GFP vector (Fig. 4a) to investigate TSWV NSm cell-to-cell movement (Feng *et al*., 2016). The expressed mCherry-HDEL binds ER membrane in the initial cells but NSm-eGFP traffics between cells. To investigate whether the Sw-5b-induced host defense can affect TSWV NSm cell-to-cell movement, we co-expressed mCherry-HDEL, NSm-GFP, and Sw-5b or mCherry-HDEL, NSm-GFP, and EV in *N. benthamiana* leaves through agro-infiltration. Under the fluorescence microscope, both NSm-GFP and mCherry-HDEL were found in single cells in the presence of Sw-5b. In the presence of EV, however, NSm-GFP moved into multiple cells, while mCherry-HDEL accumulated in the initial cells (Fig. 4b, upper two panels). The result suggested that Sw-5b elicited a defense that strongly inhibited cell-to-cell movement of viral NSm.

**Fig. 4.**
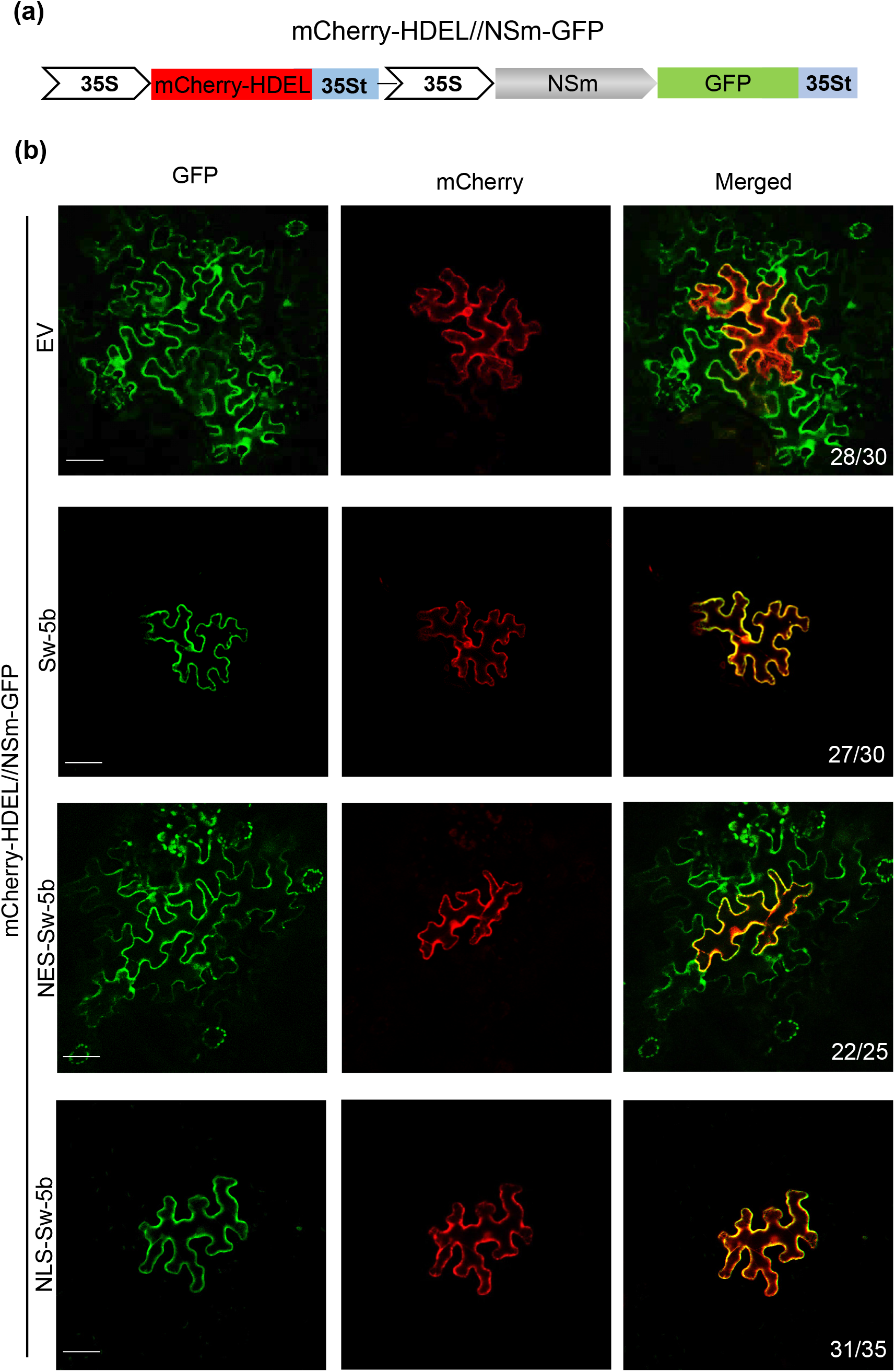
Effect of subcellular localization of Sw-5b on cell-to-cell movement of NSm in leaf epidermis of *N. benthamiana.* (a) Schematic diagram of the binary construct to co-express mCherry-HDEL and NSm-GFP. (b) Cell-to-cell movement analysis of NSm-GFP in *N. benthamiana* leaves co-expressing p2300S empty vector (EV), Sw-5b, NES-Sw-5b, NLS-Sw-5b, or nls-Sw-5b with the construct harboring both mCherry-HDEL and NSm-GFP. *Agrobacterium* containing the construct to co-express mCherry-HDEL and NSm-GFP was diluted 500 times for expression in a single epidermal cell. All other *Agrobacterium* were infiltrated at the concentration of OD600 = 0.2. Bar, 50 μm.

To make sure this inhibition to viral NSm cell-to-cell movement is not caused by overexpression of Sw-5b, we also used the NSm^T120N^ mutant, from the resistance - breaking (RB) TSWV isolates, which cannot be recognized by Sw-5b (Zhao et al., 2016). The assays showed that in the presence of either Sw-5b or EV, NSm^T120N^-GFP moved into multiple cells, while mCherry-HDEL retained in the initial cells (Fig S3a).

### Sw-5b in the nucleus but not in the cytoplasm triggers a defense against NSm cell-to-cell movement

To determine the effects of the cytoplasmic and nuclear Sw-5b on host defense against TSWV NSm intercellular movement, we co-expressed mCherry-HDEL and NSm-GFP with NES-Sw-5b, nes-Sw-5b, NLS-Sw-5b, or nls-Sw-5b in *N. benthamiana* leaves via agro-infiltration. The results showed that in the presence of NLS-Sw-5b, the cell-to-cell movement of NSm-GFP was inhibited (Fig. 4b). Similar results were also obtained in the leaves co-expressing mCherry-HDEL and NSm-GFP with nls-YFP-Sw-5b or nes-YFP-Sw-5b (Fig S3b). In the presence of NES-YFP-Sw-5b, however, NSm-GFP did move into surrounding cells. (Fig. 4b). This finding indicates that the Sw-5b in the nucleus but not in the cytoplasm induced a host defense that inhibited TSWV NSm cell-to-cell movement.

### Nuclear Sw-5b confers host immunity to TSWV systemic infection

To dissect the host immunity induced by the cytoplasmic and the nuclear Sw-5b, we generated transgenic *N. benthamiana* lines expressing YFP-Sw-5b, NES-YFP-Sw-5b, nes-YFP-Sw-5b, NLS-YFP-Sw-5b, and nls-YFP-Sw-5b, respectively (Tables S1 and S2). After inoculation of these transgenic lines with TSWV-YN isolate, the EV (control) transgenic plants developed typical viral symptoms including stunt, leaf curl and mosaic at 7 to 15 days post inoculation (dpi). The NES-YFP-Sw-5b transgenic plants developed a strong HR trailing in the systemic leaves by 7 to 15 days post inoculation (dpi) (Fig. 5a and Fig. S4a), suggesting that NES-YFP-Sw-5b transgenic plant did not block TSWV systemic infection and caused virus infection-related systemic HR. In contrast, no systemic virus infection symptoms were observed in the YFP-Sw-5b and the nes-YFP-Sw-5b transgenic plants. The RT-PCR agreed with the symptom observation results and showed that TSWV-YN genomic RNA was accumulated in the systemic leaves of the TSWV-YN-inoculated NES-YFP-Sw-5b or the EV transgenic plants, but not in the systemic leaves of the TSWV-YN-inoculated YFP-Sw-5b or nes-YFP-Sw-5b transgenic plants (Fig. 5c and Fig. S4b). Also in this study, the TSWV-YN-inoculated NLS-YFP-Sw-5b or nls-YFP-Sw-5b transgenic plants did not show virus like symptoms in their systemic leaves by 7-15 dpi (Fig. 5a, and Fig. S4a). The RT-PCR result confirmed that TSWV-N genomic RNA had not accumulated in the systemic leaves of the NLS-YFP-Sw-5b or the nls-YFP-Sw-5b transgenic plants (Fig. 5c, and Fig. S4b), indicating that the nuclear Sw-5b is responsible for the host immunity against TSWV systemic infection.

**Fig. 5.**
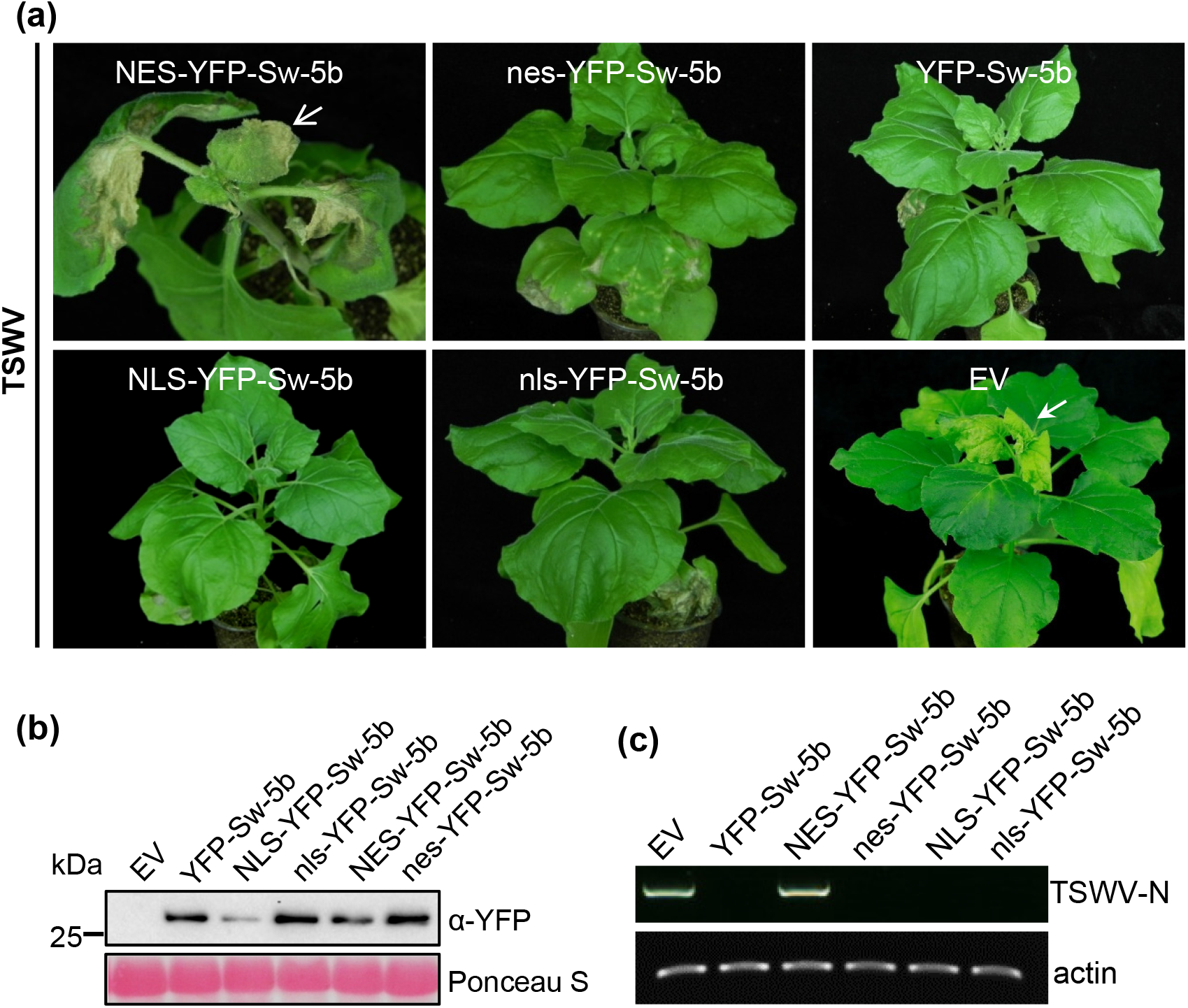
Analysis of cytoplasm- and nucleus-targeted Sw-5b-mediated host immunity to TSWV systemic infection. (a) TSWV systemic infection in transgenic *N. benthamiana* plants expressing NES-YFP-Sw-5b, nes-YFP-Sw-5b, NLS-YFP-Sw-5b, nls-YFP-Sw-5b, YFP-Sw-5b or p2300S empty vector (EV) driven by 35S promoter. TSWV-inoculated plants were photographed at 15 dpi. White arrow indicates the systemic leaves showing HR trailing. White arrowhead indicates the systemic leaves showing mosaic. (b) Immunoblot analysis of NES-YFP-Sw-5b, nes-YFP-Sw-5b, NLS-YFP-Sw-5b, nls-YFP-Sw-5b and YFP-Sw-5b expressions in different transgenic *N. benthamiana* plants. EV plants transformed with an empty vector and were used as a negative control. (c) RT-PCR analysis of TSWV accumulation in the systemic leaves of different transgenic *N. benthamiana* plants at 15 dpi.

### The cytoplasmic and the nuclear Sw-5b act synergistically to confer a strong immunity to TSWV infection in *N. benthamiana*

To investigate whether cytoplasm-targeted and nucleus-targeted Sw-5b have joint effects on the defense against TSWV infection, we constructed a M(-)opt-pSR(+)eGFP vector by inserting a cassette expressing optimized TSWV M genomic sequence (Feng *et al*., 2020) into the pSR(+)eGFP mini-replicon to express NSm, N, and eGFP simultaneously in the same cells (Fig. 6a). The construct M(-)opt-pSR(+)eGFP couples the functions for both viral replication and viral cell-to-cell movement, mimicking the virus infection in plant leaves. After co-expressing this vector, the L(+)opt and the EV in *N. benthamiana* leaves through agro-infiltration, the eGFP fluorescence was observed in many cells, due to the presence of the NSm movement protein and the RdRpopt (Fig. 6b, upper left image). When M(-)opt-SR(+)eGFP, L(+)opt and Sw-5b were co-expressed in *N. benthamiana* leaves, the eGFP fluorescence was hardly detected and some were observed only in single leaf cells (Fig. 6b upper right image, Fig. S5a and b). When M(-)opt-SR(+)eGFP, L(+)opt and NES-Sw-5b were co-expressed in *N. benthamiana* leaves, the eGFP fluorescence was observed in clusters of a few cells (Fig. 6b, Fig. S5a), indicating that limited cell-to-cell movement had occurred in these leaves (Fig. S5b). When M(-)opt-SR(+)eGFP, L(+)opt and NLS-Sw-5b were co-expressed in leaves, a few of eGFP fluorescence were detected but they were in single cells only. When leaves co-expressing M(-)opt-SR(+)eGFP, L(+)opt and NES-Sw-5b + NLS-Sw-5b, the eGFP fluorescence was also hardly detected and some were observed only in single leaf cells. Western blot results showed that more eGFP had accumulated in the leaves co-expressing M(-)opt-SR(+)eGFP, L(+)opt, and EV, followed by the leaves co-expressing M(-)opt-SR(+)eGFP, L(+)opt, and NLS-Sw-5b, and then the leaves co-expressing M(-)opt-SR(+)eGFP, L(+)opt, and NES-Sw-5b. Much less eGFP had accumulated in the leaves co-expressing M(-)opt-SR(+)eGFP, L(+)opt, and NLS-Sw-5b + NES-Sw-5b, and in the leaves co-expressing M(-)opt-SR(+)eGFP, L(+)opt, and Sw-5b (Fig. 6c and d). The accumulation of eGFP was lower in the leaves co-expressing NES-Sw-5b + NLS-Sw-5b than that in the leaves co-expressing NES-Sw-5b or NLS-Sw-5b (Fig. 6c and d), indicating that NES-Sw-5b and NLS-Sw-5b have additive role in mediating host immunity against different TSWV infection steps.

**Fig. 6.**
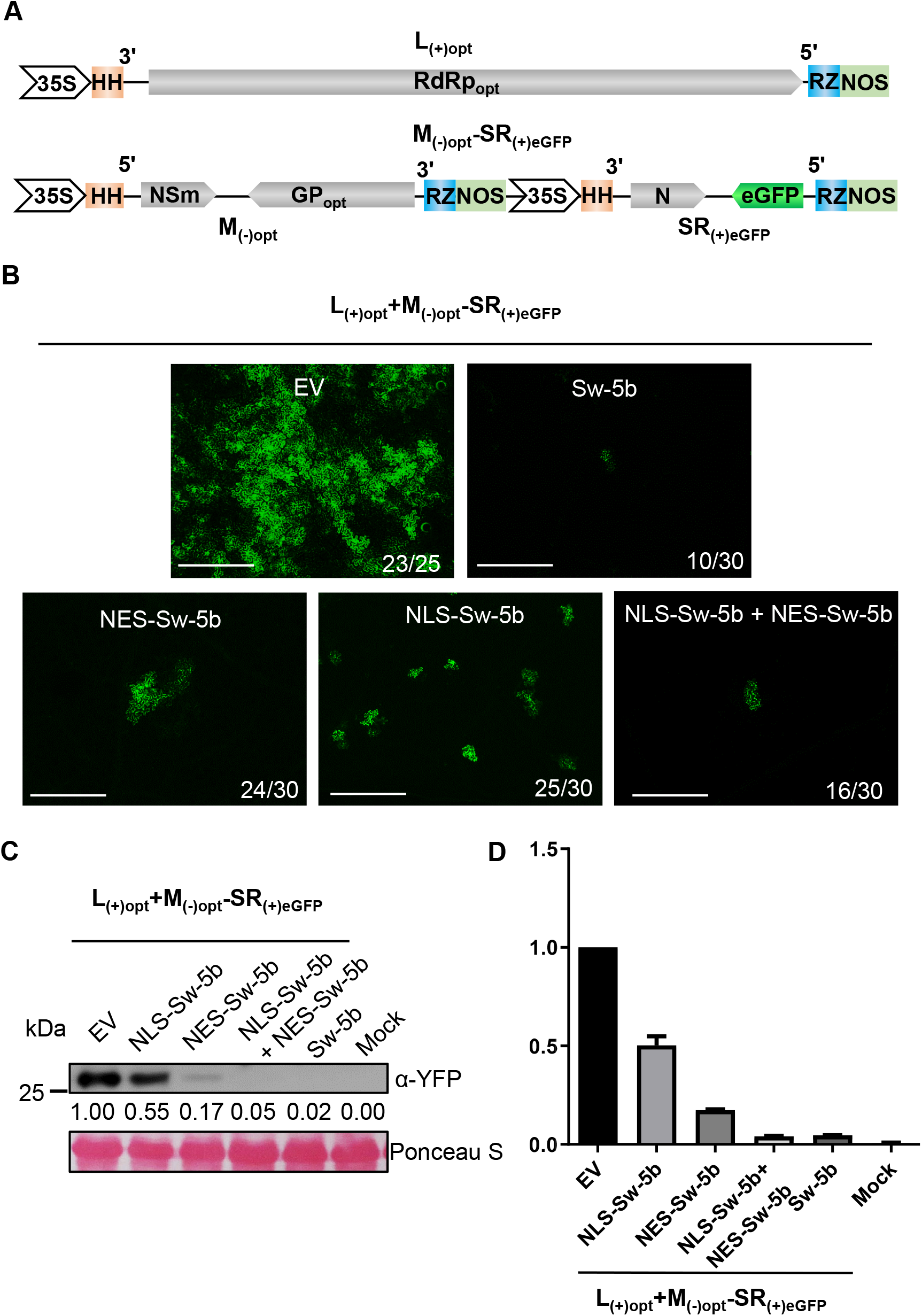
Joint effects of cytoplasm- and nucleus-targeted Sw-5b on defenses against tospovirus infection in *Nicotiana benthamiana* leaves. (a) Schematic representation of binary constructs to express TSWV SR(+)eGFP-M(-)op and TSWV L RNA segment containing an optimized RdRp. 35S: a double 35S promoter; HH: hammerhead ribozyme; RZ: hepatitis delta virus (HDV) ribozyme; NOS: nopaline synthase terminator. (b) Accumulation of eGFP fluorescence in *N. benthamiana* leaves co-expressing p2300S empty vector (EV), Sw-5b, NES-Sw-5b, NLS-Sw-5b, or NES-Sw-5b+NLS-Sw-5b with TSWV SR(+)eGFP-M(-)op at 4 days post infiltration (dpi) viewed with a fluorescence microscope. Bar represents 400 μm. (c) Immunoblot analysis of expression of eGFP proteins in leaves shown in panel (b) using specific antibodies against YFP. Ponceau S staining of rubisco large subunit is shown for protein loading control. (d) Quantification of eGFP proteins in leaves shown in panel (c).

### The Sw-5b NB-ARC-LRR control its cytoplasm localization whereas the extended N-terminal SD domain is crucial for targeting Sw-5b into nucleus, and for inducing host systemic immunity

Sw-5b has an extended N-terminal SD domain, a CC domain, a NB-ARC domain, and a C-terminal LRR domain (Chen *et al*., 2016). To determine which domain(s) of Sw-5b is/are responsible for nucleoplasm/nucleolus targeting and for plant immunity, we tested these domains using various deletion mutants and YFP fusion proteins (Fig. 7a). We reported previously that the Sw-5b NB-ARC-LRR region was able to induce HR cell death in plant in the presence of NSm (Chen *et al*., 2016). In this study, we fused YFP to the N-terminus of NB-ARC-LRR (Fig. 7a). Transient expression of YFP-NB-ARC-LRR (112 kDa) in *N. benthamiana* leaf cells resulted in a localization of the fusion protein in cytoplasm exclusively (Fig. 7b).

**Fig. 7.**
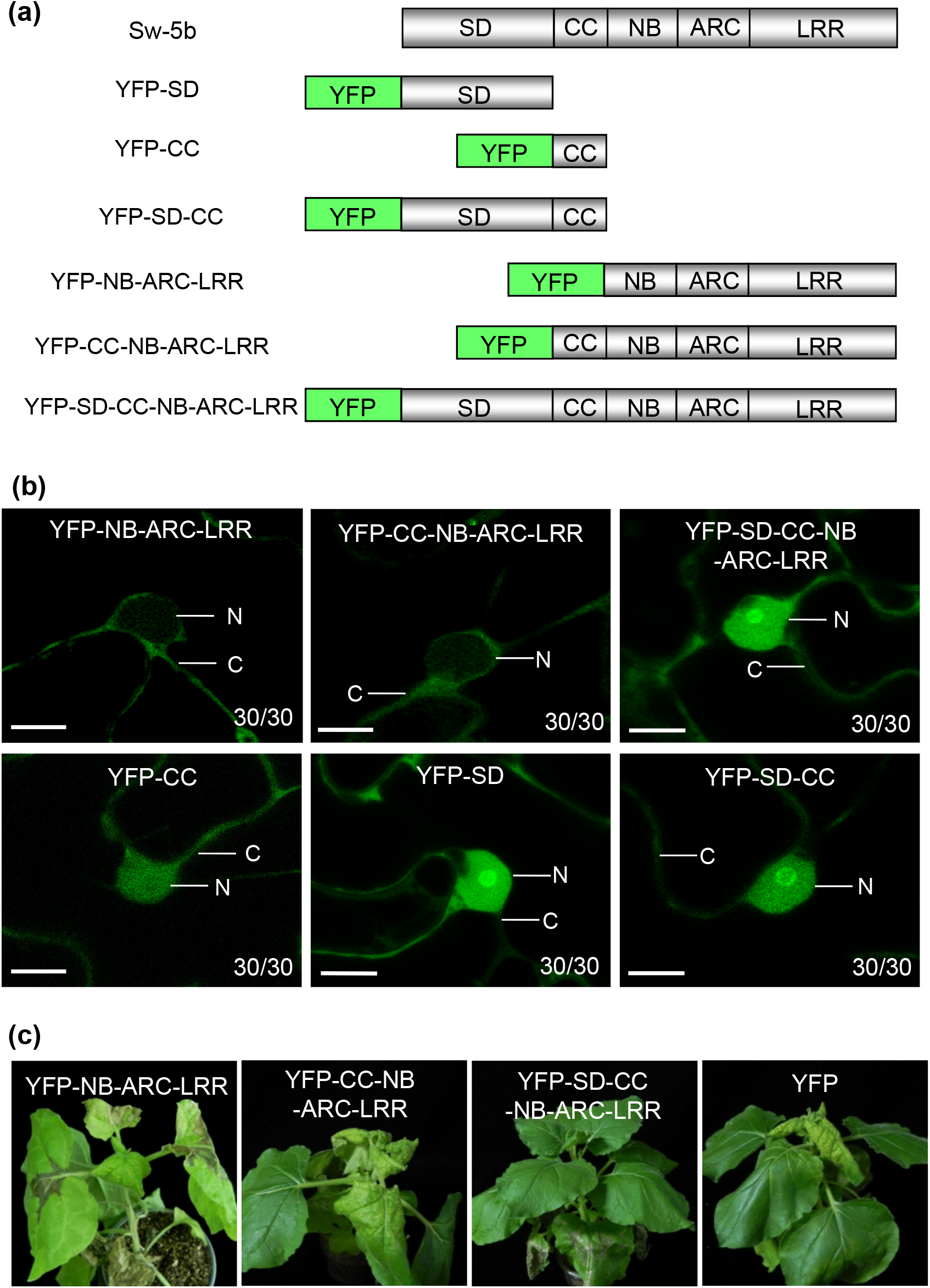
Functional analysis and subcellular localization patterns of individual or combined Sw-5b domains. (a) Schematic diagrams showing a full length Sw-5b or Sw-5b domains fused with YFP. (b) Confocal images of *N. benthamiana* leaf epidermal cells expressing various YFP fusions. Images of the cells were taken at 24 hpi. N nucleus, Nu nucleolus, and C cytoplasm. Bar = 10 µm. (c) TSWV-inoculated transgenic *N. benthamiana* plants expressing these various YFP fusions and photographed at 15 dpi.

A previous study had shown that the CC domain of potato NLR receptor Rx was required for targeting this protein to nucleus (Slootweg *et al*., 2010). To determine the function of Sw-5b CC domain in intracellular trafficking, we inserted the CC domain between the YFP and NB-ARC-LRR to generate an YFP-CC-NB-ARC-LRR construct or fused the CC domain to YFP to produce an YFP-CC construct. Transient expression of these two fusion proteins individually in *N. benthamiana* leaves, and examined the leaves under a confocal microscope, we determined that the YFP-CC fusion protein accumulated in both cytoplasm and nucleus of the cells while the YFP-CC-NB-ARC-LRR fusion protein was in the cytoplasm only (Fig. 7b). This result indicated that addition of the CC domain to YFP-NB-ARC-LRR was not sufficient to traffic the fusion protein into the nucleus.

An extended N-terminal SD domain is known to be present at the upstream of the Sw-5b CC domain. In this study, we first generated an YFP-SD and an YFP-SD-CC constructs, and transiently expressed them individually in *N. benthamiana* leaf cells. Confocal Microscopy showed that both YFP-SD and YFP-SD-CC fusion proteins accumulated in the cytoplasm and nucleus (Fig. 7b). We then inserted a SD between the YFP and CC-NB-ARC-LRR to produce an YFP-SD-CC-NB-ARC-LRR construction. Transient expression of this construct in *N. benthamiana* leaf cells showed that this fusion protein accumulated in the cytoplasm and nucleus (Fig. 7b).

We next generated stable transgenic *N. benthamiana* plants expressing YFP-NB-ARC-LRR, YFP-CC-NB-ARC-LRR and YFP-SD-CC-NB-ARC-LRR. Upon inoculation transgenic *N. benthamiana* plants expressing YFP-NB-ARC-LRR with TSWV, large HR foci were observed in the TSWV-inoculated leaves and later, HR trailing was seen in the systemic leaves of most assayed plants (Fig. 7c; Table S2 and S3). RT-PCR results confirmed the presence of TSWV genomic RNA in these systemic leaves (Fig. S6b). We also inoculate *N. benthamiana* plants expressing YFP-CC-NB-ARC-LRR with TSWV. By 7 dpi, no systemic resistance to TSWV infection was observed in these plants (Fig. 6a and B, Table S4). RT-PCR results showed that systemic infection of TSWV did occur in the TSWV-inoculated YFP-CC-NB-ARC-LRR transgenic plants (Fig. S6b). In contrast, transgenic plants expressing YFP-SD-CC-NB-ARC-LRR fusion exhibited a systemic immunity to TSWV infection (Fig. 7c, Fig. S6b).

These data indicated that Sw-5b NB-ARC-LRR control its cytoplasm localization, CC domain of Sw-5b alone was not sufficient to transport the NB-ARC-LRR into nucleus and the extended SD domain is required for targeting Sw-5b to the nucleus, and for inducing host immunity.

### The extended SD domain interacted with *importin α1*, *α2* and *β*

To identify the cellular machinery needed for transporting Sw-5b into nucleus, we co-expressed YFP-SD and YFP-Sw-5b in *N. benthamiana* leaves followed by a co-immunoprecipitation (co-IP) and Mass Spectrometry. The results identified *N. benthamiana* importin α as one of candidate proteins interacted with YFP-SD and YFP-Sw-5b (Table S5 and S6). The co-IP and Mass Spectrometry also identified nuclear pore complex protein TPRb and nuclear pore complex protein Nup160a interacted with YFP-Sw-5b (Table S6). *Importins* play important roles in translocating proteins from cytoplasm into nucleus (Kanneganti *et al*., 2007). We used BiFC analysis to confirm the interaction between YFP-SD with *N. benthamiana* importin homologs α1, α2 and *β*. The result showed that co-expression of cYFP-SD with nYFP-IMP α1, nYFP-IMP α2 or nYFP-IMP *β* produced a strong YFP fluorescence signal in nucleus. Co-expression of cYFP-Sw-5b with nYFP-IMP α1, nYFP-IMP α2 or nYFP-IMP *β* also detected a strong YFP fluorescence signal in nucleus (Fig. S7). In contrast, co-expression of controls cYFP-SD and nYFP, cYFP-Sw-5b and nYFP, cYFP and nYFP-IMP α1, cYFP and nYFP-IMP α2 or cYFP and nYFP-IMP *β* did not show fluorescence signal in *N. benthamiana* leaf cells (Fig. S7).

### Silencing *importin α1*, *α2* and *β* expression abolished Sw-5b nucleus accumulation and host resistance to TSWV systemic infection

To determine the functions of importin α1, α2 and β in Sw-5b nucleus localization, we silenced *importin α1, α2, β, α1* and *α2,* and *α1* and *α2* and *β* expressions, respectively, in *N. benthamiana* leaves using a tobacco rattle virus (TRV)-based virus-induced gene silencing (VIGS) vector, and then transiently expressed YFP-Sw-5b in these plants. Analyses of these plants through RT-PCR using gene specific primers showed that silencing of these *importin* genes in *N. benthamiana* leaves were successful (Fig. S8a). However, silencing individual *importin* gene or both *importin α1* and *α2* was not enough to block the nucleus accumulation of YFP-Sw-5b (Fig. 8a). In contrast, after *importin α1*, *α2* and *β* were all silenced through VIGS, the nucleus accumulation of YFP-Sw-5b was inhibited (Fig. 8a, the middle image in the bottom panel).

**Fig. 8.**
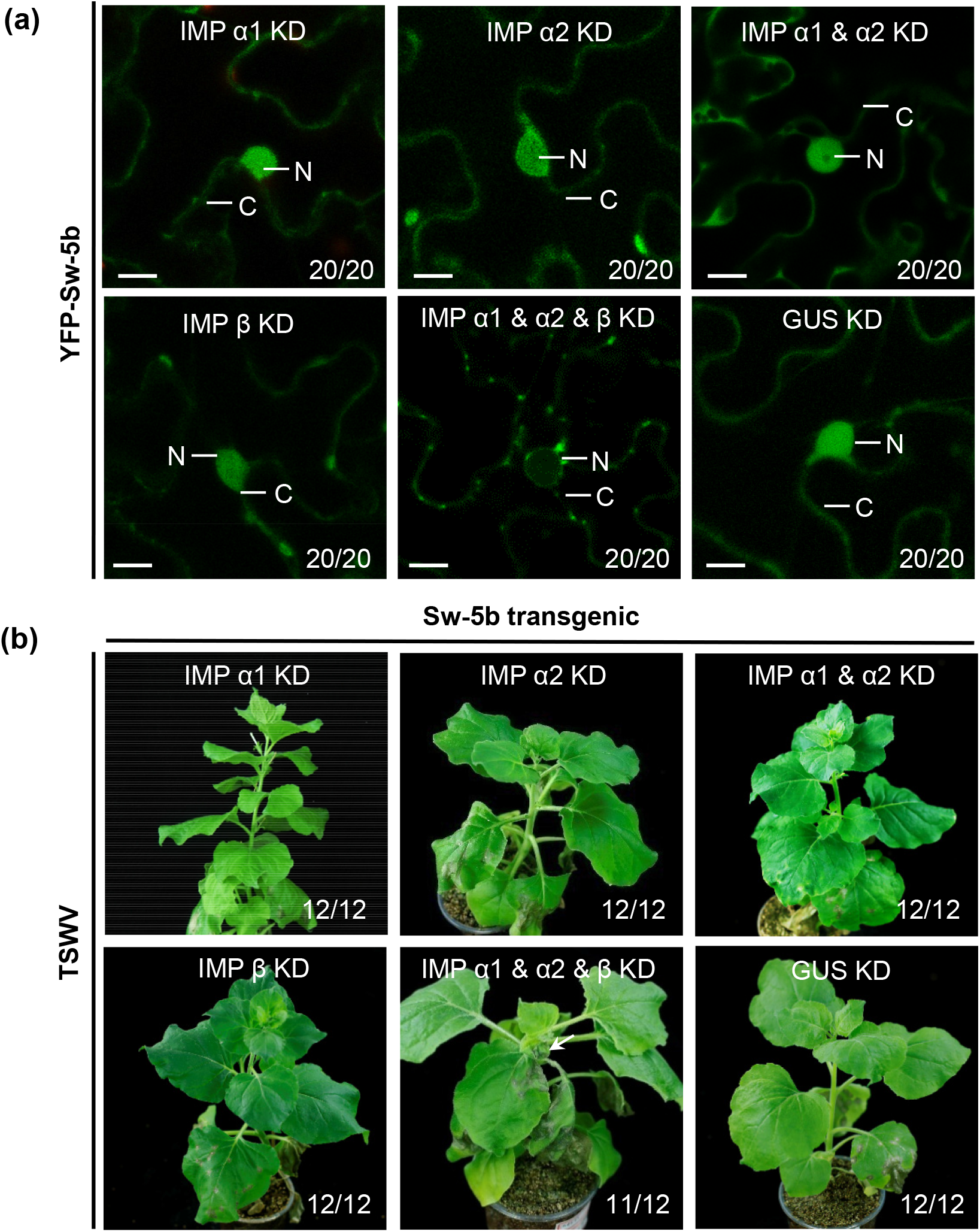
Roles of *importins α* and *β* in YFP-Sw-5b nucleus targeting and Sw-5b-mediated immunity to TSWV systemic infection. (a) Transient expression of YFP-Sw-5b in *N. benthamiana* leaf epidermal cells silenced for *importin α1* (IMP α1 KD)*, importinα2* (IMP α2 KD), *importin α1* and *α2* (IMP α1 & α2 KD), *importin β* (IMP β KD) or *importin α1* and *α2* and *β* (IMP α1 & α2 & β KD) through VIGS. Images of the cells were captured using a confocal microscope at 26 hpi. N nucleus, C cytoplasm. Bar = 10 μm. (b) YFP-Sw-5b transgenic *N. benthamiana* plants were silenced for *importin α1, importin α2*, *importin α1* and *α2*, *importin β* or *importin α1* and *α2* and *β* expression through VIGS followed by inoculation with TSWV. TSWV-inoculated YFP-Sw-5b transgenic *N. benthamiana* plants were photographed at 15 dpi. White arrowhead indicates HR trailing in systemic leaves.

To investigate the effects of nuclear import defected Sw-5b on host immunity to TSWV systemic infection, we silenced these *importin* genes in the Sw-5b transgenic *N. benthamiana* plants as described above, and then inoculated them with TSWV. The results showed that the plants silenced for *importin α1*, *α2*, and *β* gene, individually, did not show TSWV systemic infection (Fig. 8b and Fig. S8b). In addition, the transgenic plants silenced for both *importin α1* and *α2* genes also did not show TSWV systemic infection (Fig. 8b and Fig. S8b). In contrast, after silencing *importin α1*, *α2* and *β* together, the plants developed clear TSWV symptoms in systemic leaves followed by HR (Fig. 8b, white arrow and Fig. S8b), indicating that the nucleus accumulation of Sw-5b is indispensable for the induction of host immunity against TSWV systemic infection.

## Discussion

In this report, we provide evidence to show that the cytoplasm-accumulated and the nucleus-accumulated Sw-5b, a tomato immune receptor, play different roles in inducing host defense against TSWV infection in plant. The cytoplasmic Sw-5b functions to induce a strong cell death response to inhibit TSWV replication. This host response is, however, insufficient to block virus intercellular and long-distance movement. The nuclear-localized Sw-5b triggers a host defense that weakly inhibit viral replication but strongly inhibit tospovirus intercellular and systemic movement. These findings suggest that tomato Sw-5b NLR induces different types of defense responses by cytoplasm and nucleus partitioning to combat virus at different infection steps. Furthermore, the cytoplasmic and the nuclear Sw-5b act synergistically to confer a strong host immunity to TSWV infection in plant. We also demonstrated that the extra SD domain functioned as a critical intracellular translocation modulator, allowing Sw-5b receptor to translocate from cytoplasm to nucleus to trigger the immunity. The Sw-5b NB-LRR controls its cytoplasm localization. Unlike Rx CC domain, Sw-5b CC domain is not sufficient to translocate NB-LRR into nucleus. Strikingly, the SD is crucial for Sw-5b to translocate from cytoplasm for nucleus. This SD-mediated receptor translocation is dependent on importins α and β.

Successful virus infection in plant requires several steps including viral replication in the initially infected cells followed by cell-to-cell and long-distance movement (Heinlein, 2015; Wang, 2015). After entering into plant cells, virus first encode multiple proteins needed for its replication. Once the initial replication is established, virus will encode specific protein(s), known as movement proteins (MPs), to traffic viral genome or virions into adjacent cells through plasmodesmata in cell walls, and then long-distantly into other parts of the plant to cause a systemic infection (Rao, 2002; Lucas, 2006; Taliansky *et al*., 2008). To date, multiple plant NLRs, conferring host resistance against plant viruses have been identified (Soosaar *et al*., 2005; Meier *et al*., 2019), but how these plant NLRs induce host resistance against virus infection remain largely unknown.

In this study, we have determined that the forced cytoplasm accumulation of Sw-5b can induce a stronger cell death than that caused by the accumulation of Sw-5b in both cytoplasm and nucleus. While, the cell death induced by the forced nucleus accumulation of Sw-5b was significantly weakened. We then analyzed Sw-5b-mediated immunity against TSWV replication using a TSWV mini-replicon system and a movement defective NSm mutant. Our results showed that the forced cytoplasm accumulation of Sw-5b can induce a strong host defense against virus replication in cells. This finding implies that cytoplasm is one of the main source of defense signaling against TSWV replication. The defense signaling generated in nucleus can only induce a weak defense against TSWV replication. Therefore, the nuclear localized Sw-5b is only partially responsible for the induction of host defense against TSWV replication. It is also possible that this nuclear localized Sw-5b-induced weak host response is caused by a trace of NLS-YFP-Sw-5b maintained in the cytoplasm that maybe below the detection limit of Confocal Microscope. It has been shown to accumulate in both cytoplasm and nucleus, and the forced cytoplasm accumulation of Barley MLA10 enhance cell death signaling (Bai *et al*., 2012). We speculate that, for both MLA10 and Sw-5b, the cytoplasm accumulation is crucial for the initiation and/or amplification of the cell death signaling. The CC and the TIR domain of several plant NLRs have been shown to trigger cell death (Swiderski *et al*., 2009; Krasileva *et al*., 2010; Bernoux *et al*., 2011; Collier *et al*., 2011; Maekawa *et al*., 2011; Bai *et al*., 2012; Chen *et al*., 2017; Wang *et al*., 2020). Analyses of the three dimensional structures of Arabidopsis ZAR1 resistosome have also shown that its CC domain can form pentamer structures that was able to target into host cell membranes, leading to ion leakage and cell death (Wang *et al*., 2019a; Wang *et al*., 2019b). We speculate that cell death likely cause the toxicity on viral replicase or other proteins associated with virus replication in cells.

Plant virus encodes specific movement protein(s) to traffic viral genome between cells and then leaves to cause systemic infection (Rao, 2002; Lucas, 2006; Taliansky *et al*., 2008). We reported previously that TSWV NSm alone can move between plant cells (Feng *et al*., 2016). In this study, we investigated the effect of the Sw-5b-mediated host defense on TSWV intercellular movement. Through this study, we have determined that after the recognition of NSm, Sw-5b receptor induced a strong reaction to block NSm intercellular trafficking. Previous reports have some indications on the role of plant NLRs in viral movement. Nevertheless it has no direct evidence showing that plant NLRs induce resistance against viral movement. Deom and colleagues had shown that the 9.4-kDa fluorescein isothiocyanate-labeled dextran was unable to move between cells in the transgenic tobacco *N* leaves expressing tobacco mosaic virus (TMV) movement protein at 24°C, an HR-permissive temperature (Deom *et al*., 1991). However, that study did not involve a TMV Avr protein. In a different report, TMV-GFP showed a limited cell-to-cell movement in leaves of tobacco cv. Sumsan NN at 33°C, an HR-nonpermissive temperature (Canto & Palukaitis, 2002). Li and colleagues found that after treatment of SMV-inoculated Jidou 7 resistant plants with a callose synthase inhibitor, the plants showed enlarged HR lesions (Li *et al*., 2012). The soybean *Rsv3* induced extreme resistance. However, after this extremely resistant soybean line was treated with a callose synthase inhibitor, the plants developed HR lesions upon SMV-G5H inoculation (Seo *et al*., 2014). These reports indicate that plant NLRs likely involves the defense against viral movement. Here we provide the direct evidence that Sw-5b NLR can induce a strong defense response to impede NSm intercellular trafficking. More importantly, we have determined that the induction of host immunity to TSWV intercellular movement requires the accumulation of Sw-5b in nucleus. Although the cytoplasmic Sw-5b can induce a strong cell death response, it cannot prevent TSWV NSm cell-to-cell movement. Consequently, we propose that nucleus is a key compartment to generate defense signaling to block TSWV cell-to-cell movement.

In this study, although the NES-YFP-Sw-5b transgenic *N. benthamiana* plants showed an HR, they were unable to stop TSWV systemic infection. We also showed that Sw-5b YFP-NB-ARC-LRR (112 kDa) accumulates in cytoplasm exclusively (Fig. 7b), however, transgenic *N. benthamiana* plants expressing YFP-NB-ARC-LRR show strong systemic HR trailing caused by TSWV infection. Based on these findings, we conclude that HR cell death alone is not sufficient to block TSWV long-distance movement. In our study, the NLS-YFP-Sw-5b transgenic plants were resistant to TSWV systemic infection. After silencing the expressions of *importin α1*, *α2* and *β* simultaneously to inhibit the nucleus accumulation of Sw-5b, however, the resistance to TSWV systemic infection was abolished. These findings indicate that the Sw-5b-mediated resistance signaling against viral systemic infection is generated in nucleus. Some plant NLRs are known to interact with specific transcription factors in nucleus upon recognition of pathogen effectors (Cui *et al*., 2015; Kapos *et al*., 2019). The immune regulator EDS1 has also been shown to accumulate in nucleus to reprogram RNA transcription (Garcia *et al*., 2010; Heidrich *et al*., 2011; Cui *et al*., 2015; Lapin *et al*., 2020). How Sw-5b regulates host immunity in nucleus requires further investigations.

Several plant immune receptors and immune regulators, including, e.g. potato Rx (Slootweg *et al*., 2010; Tameling *et al*., 2010), tobacco N (Burch-Smith *et al*., 2007; Caplan, JL *et al*., 2008), barley MLA10 (Shen *et al*., 2007), Arabidopsis RRS1-R/RPS4, and snc1 (Deslandes *et al*., 2003; Wirthmueller *et al*., 2007; Cheng *et al*., 2009), as well as Arabidopsis NPR1 (Katagiri & Tsuda, 2010), and EDS1 (Lapin *et al*., 2020) have been found to be nucleocytoplasmic. For some of them, nuclear accumulation of NLRs are required for the induction of plant immunity to pathogen attacks. Moreover, the MLA10-YFP-NES fusion was found to induce a strong cell death response, but not a strong host resistance to powdery mildew fungus infection. In contrast, the MLA10-YFP-NLS fusion inhibited its activity to induce a cell death response, but caused a host immunity to this pathogen (Bai *et al*., 2012). In many plant-pathogen interactions, cell death responses can be uncoupled from disease resistance (Bendahmane *et al*., 1999; Gassmann, 2005; Coll *et al*., 2010; Heidrich *et al*., 2011). This separation raises questions about how host resistance prevents pathogen invasion and what are the roles of cell death during pathogen infection. It is unclear whether the MLA10-YFP-NES-induced cell death has some inhibitory effects on powdery mildew fungus infection. In this study, we determined that cytoplasm- and nuclear-accumulation of Sw-5b have different functions. The cytoplasm-accumulated Sw-5b induces a strong defense against virus replication, whereas the nuclear-accumulated Sw-5b induced an inhibition of virus cell-to-cell and long distance movement. Both cytoplasmic and nuclear Sw-5b are needed to confer a synergistic and full defense against tospovirus infection.

We have also determined that Sw-5b NB-ARC-LRR and SD domains are important to regulate the proper subcellular localization of Sw-5b and the proper nucleoplasmic distribution of Sw-5b is needed to elicit full immune responses to inhibit different TSWV infection steps. Sw-5b NB-ARC-LRR controls its cytoplasm localization. The CC domain of the Sw-5b is not sufficient to target the receptor into nuclear. Importantly, the extended SD of Sw-5b is absolutely required for the nucleus translocation. Because non-canonical domains are frequently found in other NLRs and are quite diversified, our findings have broad implications to investigate the potential new functions of non-canonical domains that integrated in the plant NLRs to regulate the plant immunity against pathogen invasions.

Through co-IP, Mass Spectrometry and BiFC analysis, we found that the extended SD domain and Sw-5b interacted with host importin machineries to translocate the Sw-5b receptor from cytoplasm into nucleus to mediate local and systemic resistance to tospovirus. Recent studies have shown that nuclearporin MOS3, MOS6 and nuclear pore complex component MOS7/Nup88 proteins played important roles in regulating Arabidopsis innate immunity (Palma *et al*., 2005; Zhang & Li, 2005; Cheng *et al*., 2009). We found that when *importin α1*, *importin α2* and *importin β* gene were all silenced through VIGS, the nucleus targeting of Sw-5b were completely blocked, and consequently, the Sw-5b-mediated systemic immunity to tospovirus infection was compromised. Importin α and β are known to form a nucleus import complex. Binding with importin β could activate importin α to form a binding surface for NLS proteins (Stewart, 2007). Hence, disruption of either importin α or importin β would block the nucleus import of NLS proteins. Because silencing *importin α* or *importin β* gene expression through VIGS did not disrupt the nuclear targeting of Sw-5b, we speculate that a non-canonical nuclear import pathway may take part in importing Sw-5b into nucleus.

Based on the above results, we have created a working model for the Sw-5b NLR-induced host resistance against TSWV replication, and intercellular and long-distance movement in plant (Fig. 9). Upon recognition of NSm in cytoplasm, Sw-5b switched from an autoinhibited state to an activated state. The activated Sw-5b accumulated in cytoplasm and also translocate into nucleus via importins α and β. The cytoplasm-accumulated and the nucleus-accumulated Sw-5b play different roles in inducing host immunity against TSWV infection. The cytoplasmic Sw-5b functions to induce a cell death response to inhibit TSWV replication, while the nuclear Sw-5b functions to induce a weak host defense against TSWV replication, but a strong defense against TSWV cell-to-cell and long-distance movement. The concerted defense signaling generated in the cytoplasm and nucleus resulted in a strong host resistance to tospovirus infection.

**Fig. 9.**
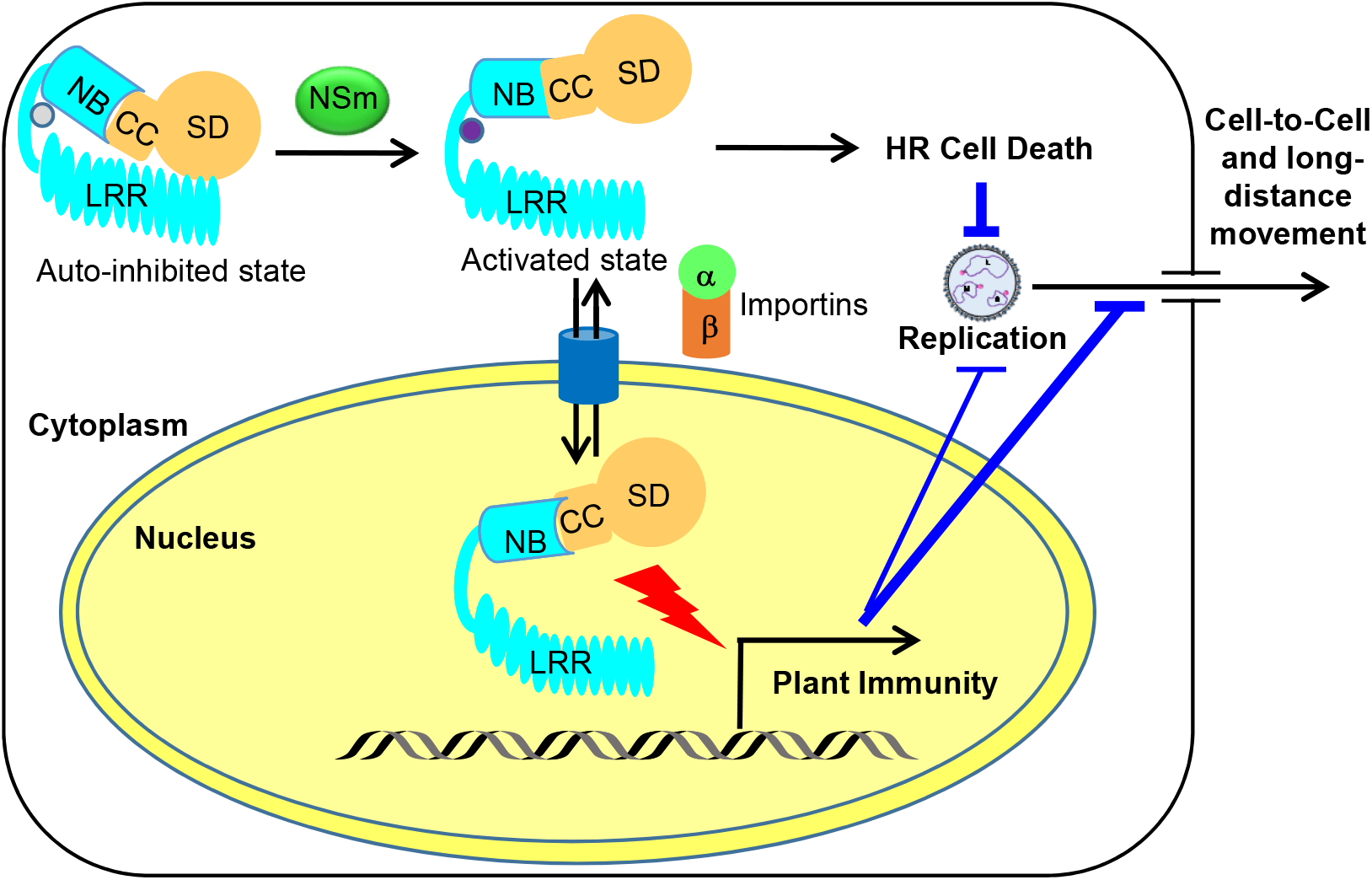
A working model for Sw-5b. Sw-5b furcates disease resistances by proper nucleocytoplasmic partition to block different infection steps of tomato spotted wilt tospovirus. Sw-5b switched from the autoinhibited state to an activated state upon recognition of NSm in the cytoplasm. Cytoplasm portion of Sw-5b induce cell death and defense that inhibit viral replication. The activated Sw-5b also translocated into nucleus via *importins α* and *β*. Nucleus-localized Sw-5b induces a defense that block viral cell-to-cell and long-distance movement. Cytoplasm- and nucleus-localized Sw-5b have additively effects on defense to inhibit viral replication, intercellular and long-distance movement during tospovirus infection.

## Acknowledgments

This work was supported by the National Natural Science Foundation of China (31630062, 31925032 and 31870143), the Fundamental Research Funds for the Central Universities (JCQY201904 and KYXK202012), Youth Science and Technology Innovation Program to XT.

## Author contributions

HC XQ, XC and XT, designed the research; HC, XQ and XC, TY, MF, JC, RC, HH, YZ, YM, DS, YX, MZ performed the experiments; HC, XSD and XT interpreted the result and wrote the paper.

## Data availability

All data produced in this study are presented in this manuscript or as the supporting files

## Supporting Information

**Fig. S1.**
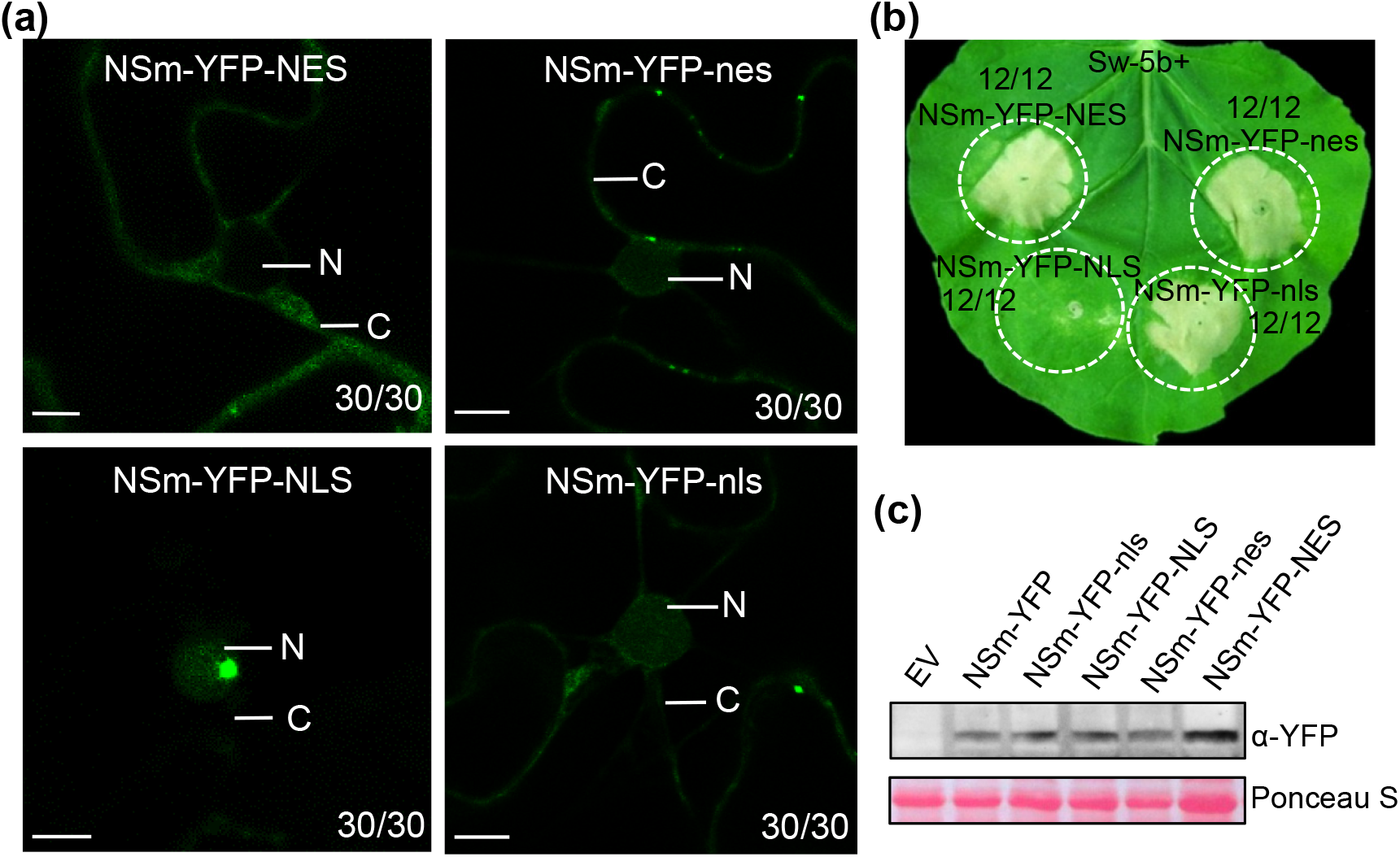
Sw-5b recognizes TSWV NSm in cytoplasm. (a) Transient expressions of NSm-YFP-NES, NSm-YFP-nes, NSm-YFP-NLS, and NSm-YFP-nls, respectively, in *N. benthamiana* leaves through agro-infiltration. Epidermal cells expressing various fusion proteins were imaged under a confocal microscope at 24 hpai. The numbers in each image indicate the number of cells showing this subcellular localization pattern and the total number of cells examined per treatment. N, nucleus; C, cytoplasm. Bar = 10 μm. (b) Various fusion proteins described in (a) were, individually, co-expressed with Sw-5b in *N. benthamiana* leaves. A representative leaf was photographed at 5 dpai. (c) Western blot analysis of various NSm fusion protein expressions in the assayed *N. benthamiana* leaves using a YFP specific antibody. Leaf areas co-expressing Sw-5b and EV were used as negative controls. The Ponceau S stained Rubisco large subunit gel was used to show sample loadings.

**Fig. S2.**
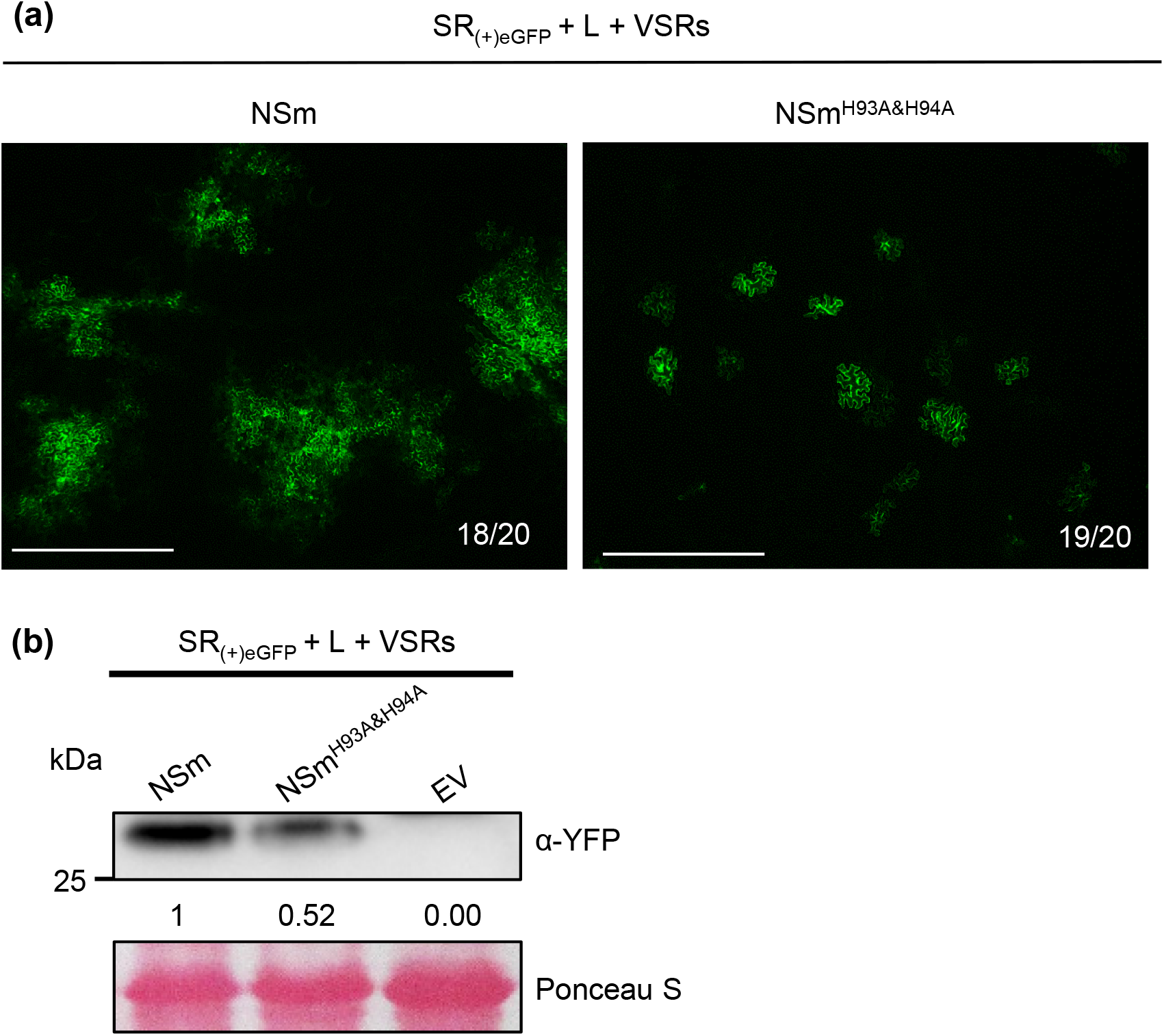
Analysis of virus replication monitoring system using a TSWV- based mini-genome replicon and a movement defective NSm mutant. (a) SR_(+)eGFP_, L_(+)opt_, VSRs and NSm or NSm^H93A&H94A^ mutant were transiently co-expressed in *N. benthamiana* leaves through agro-infiltration. The infiltrated leaves were examined and imaged under a confocal microscope at 4 dpai. The numbers in each image indicate the number of cells showing similar expression pattern and the total number of cells examined per treatment. Bar = 400 μm. (b) Western blot analysis of eGFP accumulation in assayed leaves using a YFP specific antibody. Leaves co-expressing SR_(+)eGFP_, L_(+)opt_, VSRs and EV were used as negative controls. The Ponceau S stained Rubisco large subunit gel was used to show sample loadings.

**Fig. S3.**
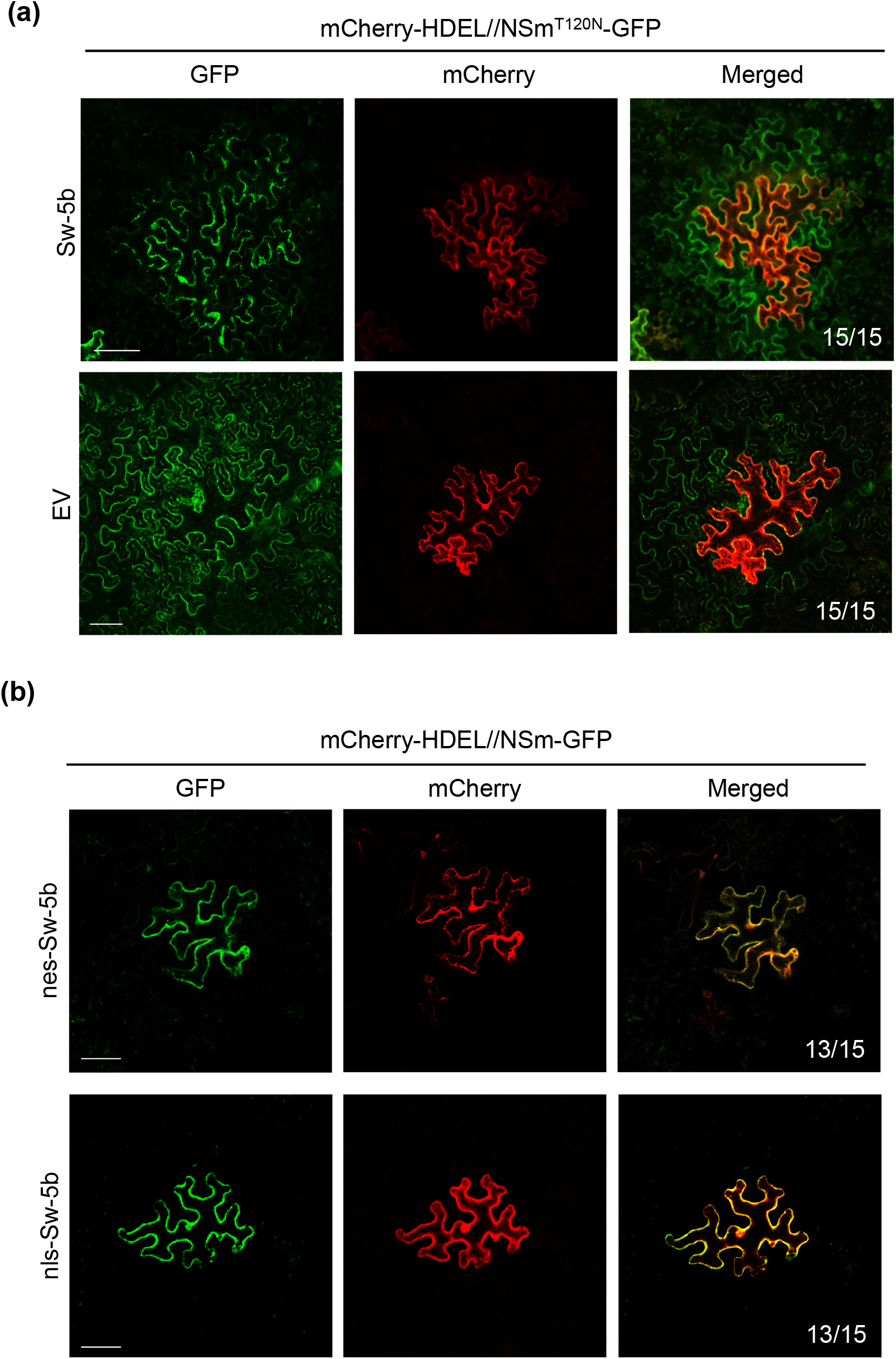
Effects of nes-Sw-5b and nls-Sw-5b on NSm-GFP cell-to-cell movement and effects of Sw-5b and EV on NSm^T120N^-GFP cell-to-cell movement. (a) Sw-5b and EV were, respectively, co-expressed with mCherry-HDEL//NSm^T120N^-GFP in *N. benthamiana* leaves through agro-infiltration. (b) nes-Sw-5b and nls-Sw-5b were, respectively, co-expressed with mCherry-HDEL//NSm-GFP in *N. benthamiana* leaves through agro-infiltration. The Agrobacterium culture carrying pmCherry-HDEL//NSm-GFP or pmCherry-HDEL//NSm^T120N^-GFP was first adjusted to OD_600_ = 0.2 and then further diluted 500 times prior to use. All other Agrobacterium cultures were adjusted to OD_600_ = 0.2 prior to use. The numbers in each image indicate the number of cells showing similar expression pattern and the total number of cells examined per treatment. Bar = 50 μm.

**Fig. S4.**
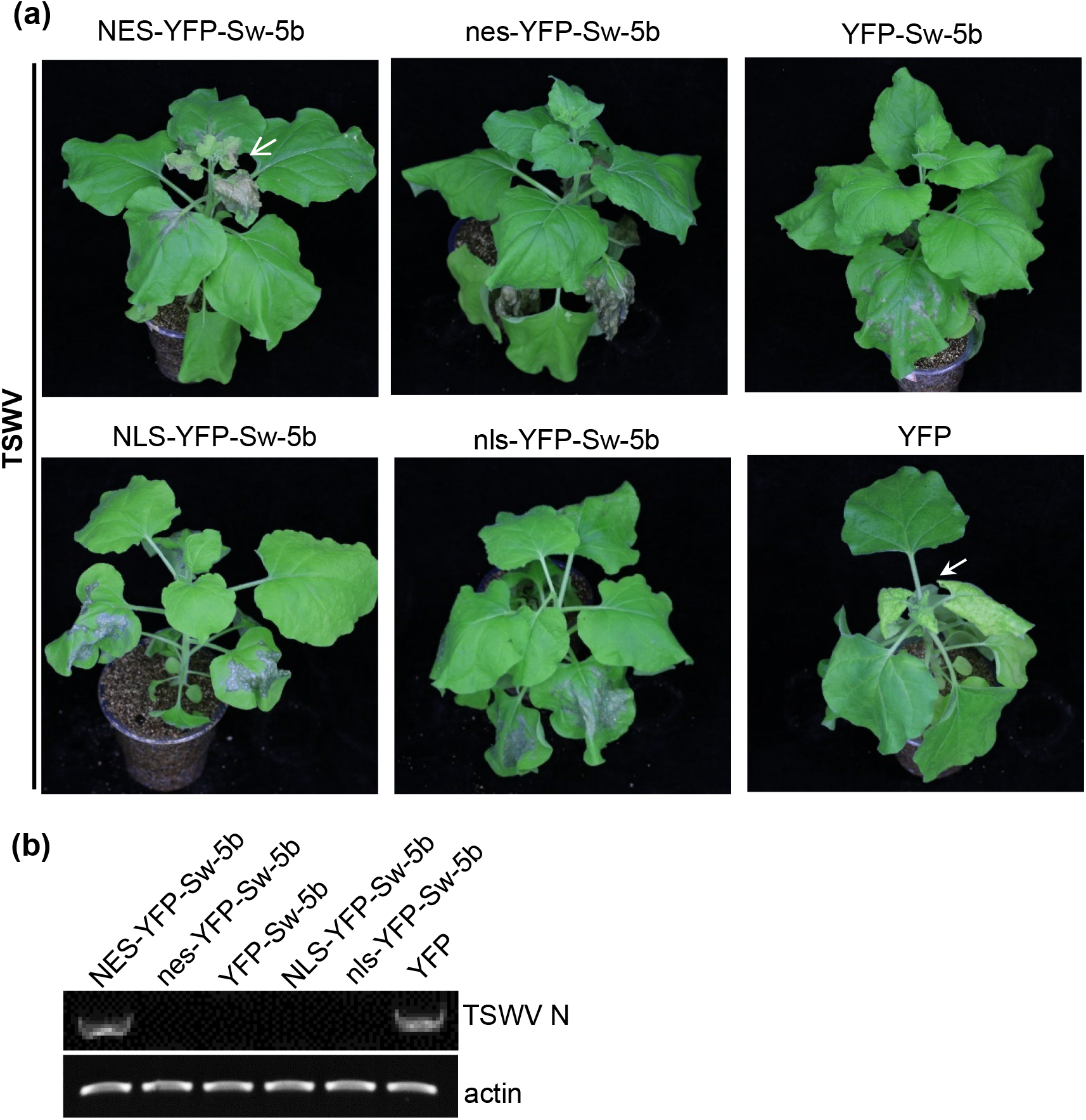
Effects of cytoplasmic and nuclear Sw-5b on host immunity to TSWV systemic infection. (a) Transgenic *N. benthamiana* lines expressing NES-YFP-Sw-5b, nes-YFP-Sw-5b, NLS-YFP-Sw-5b, nls-YFP-Sw-5b or YFP-Sw-5b, driven by the Sw-5b promoter, were used in this study. The EV transgenic plants were used as controls. The transgenic plants were inoculated with TSWV and photographed at 15 dpi. White arrow indicate the systemic leaves showing HR trailing. White arrowhead indicates the systemic leaves showing mosaic symptoms. (b) RT-PCR detection of TSWV infection in the systemic leaves of the assayed *N. benthamiana* plants at 15 dpi.

**Fig. S5.**
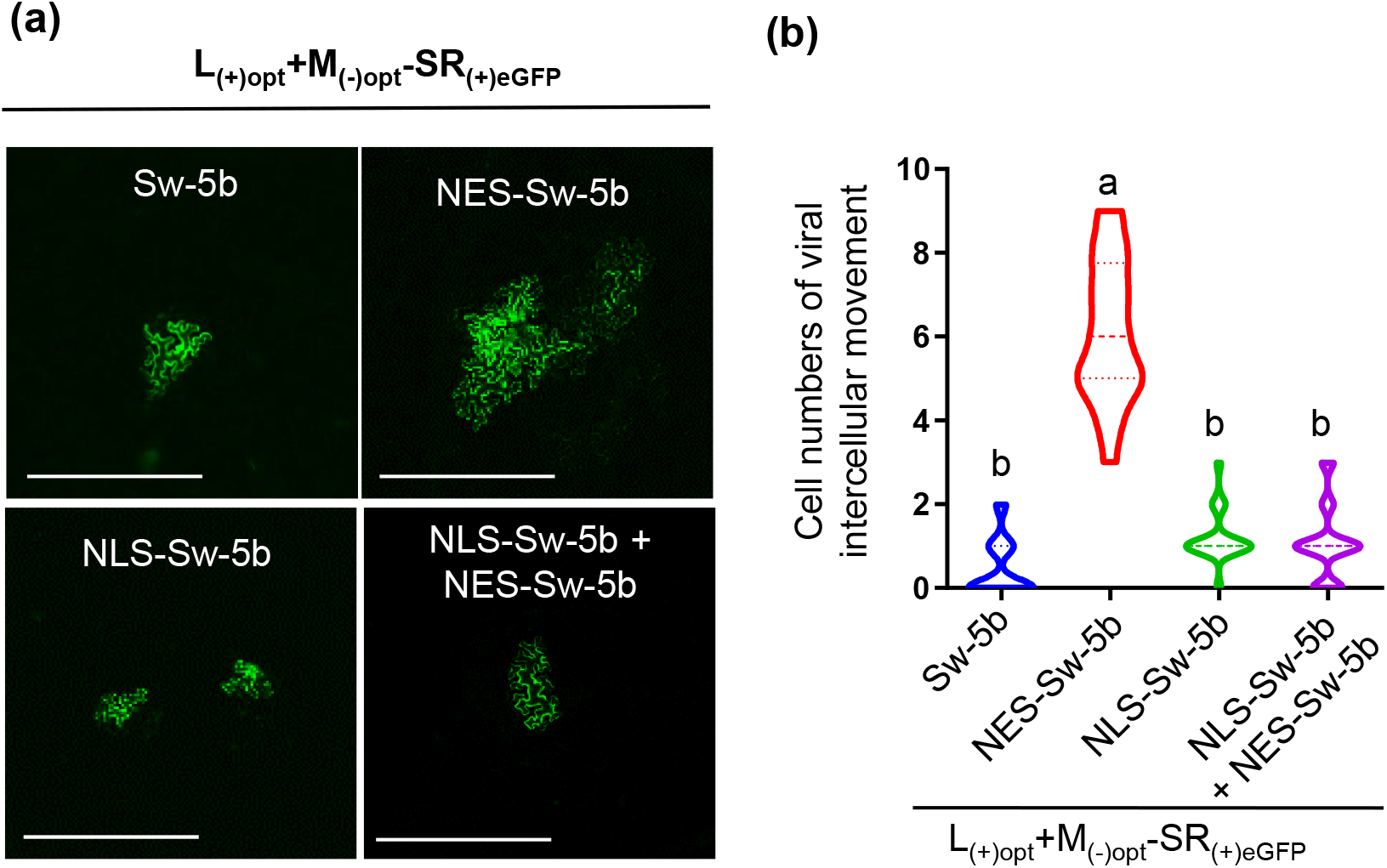
Cytoplasmic and nuclear Sw-5b activity on TSWV-GFP cell-to-cell movement in *N. benthamiana* leaves. (a) pL_(+)opt_ and pSR_(+)eGFP_-M_(-)opt_ were co-inoculated with TSWV-GFP into *Nicotiana benthamiana* leaves through agro-infiltration. The inoculated leaves were examined and imaged under a confocal microscope at 4 dpai. Bar = 400 μm. (b) Statistic analysis of TSWV-GFP cell-to-cell movement in the assayed *N. benthamiana* leaves from Figure 7B. A total of 9 assayed leaves were used for each treatment.

**Fig. S6.**
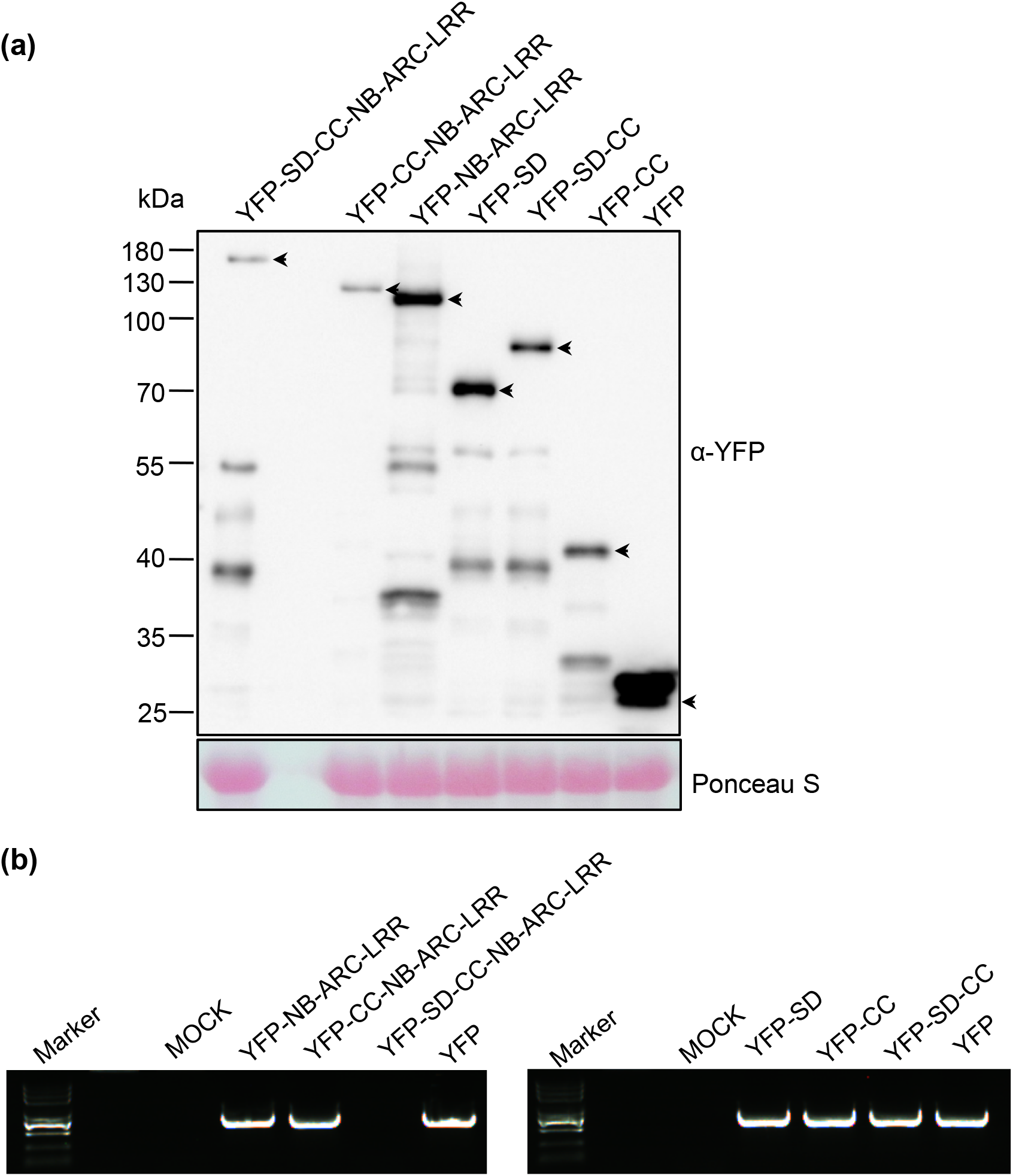
An immunoblot showing the accumulations of various YFP-tagged proteins expressed in different transgenic *N. benthamiana* plants and RT-PCR analysis of TSWV accumulation in the systemic leaves. (a) The fusion proteins were detected using an YFP specific antibody. Arrows indicate the positions of the expressed fusion proteins. Ponceau S stained Rubisco large subunits were used to estimate sample loadings. (b) RT-PCR analysis of TSWV accumulation in the systemic leaves of different transgenic *N. benthamiana* plants at 15 dpi.

**Fig. S7.**
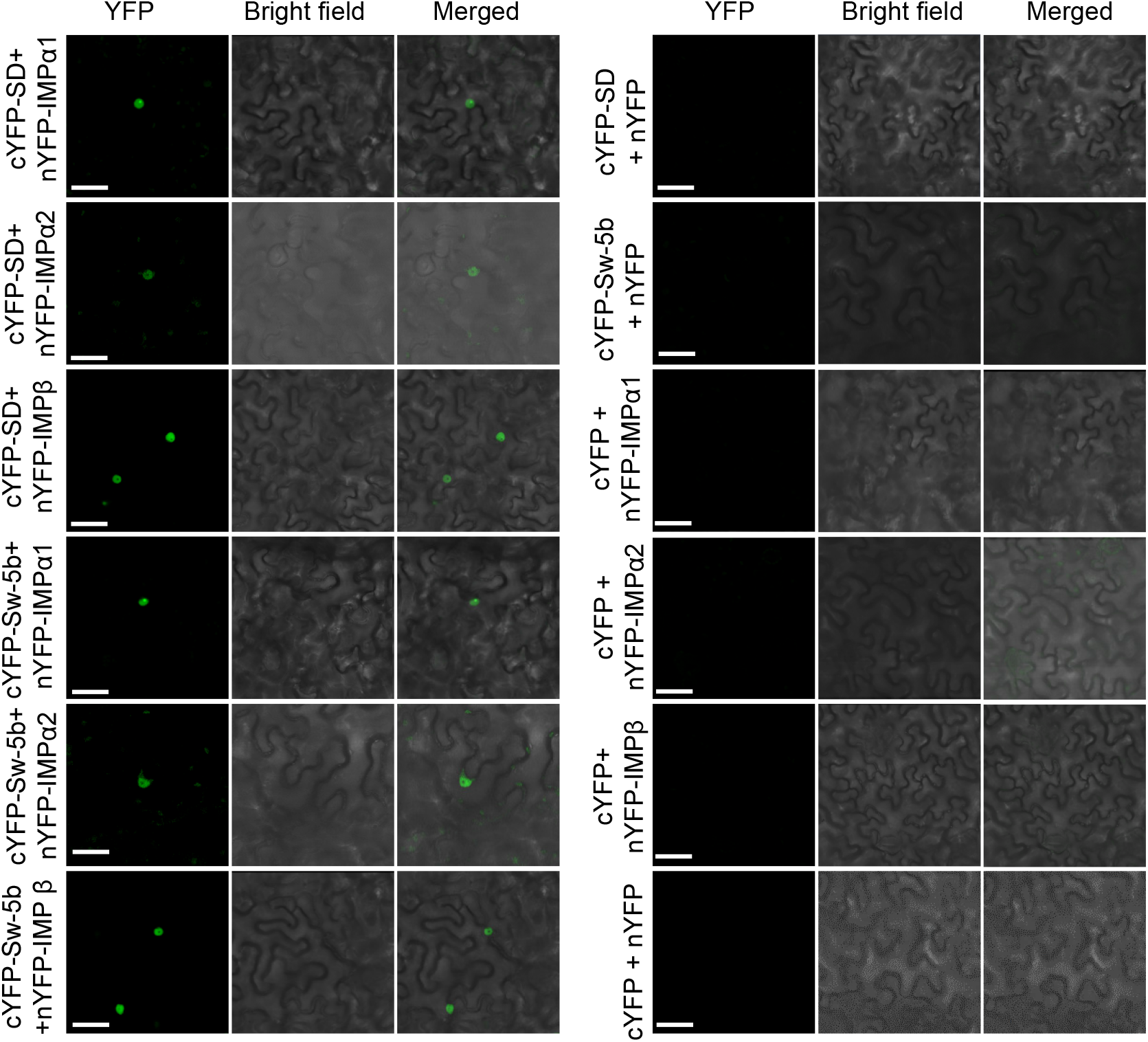
Bimolecular fluorescence complementation (BiFC) assay of cYFP-SD, cYFP-Sw-5b and nYFP-Importin α1 (nYFP-IMP α1), nYFP-Importin α2 (nYFP-IMP α2), nYFP-Importin β (nYFP-IMP β) interaction in N. benthamiana leaf epidermal cells. Confocal images were taken at 26 hour post agro-infiltration. Leaf epidermal cells co-expressing cYFP-SD and nYFP, cYFP-Sw-5b and nYFP or cYFP and nYFP-IMP α1, nYFP-IMP α2, nYFP-IMP β were used as controls. Bar = 50 μm.

**Fig. S8.**
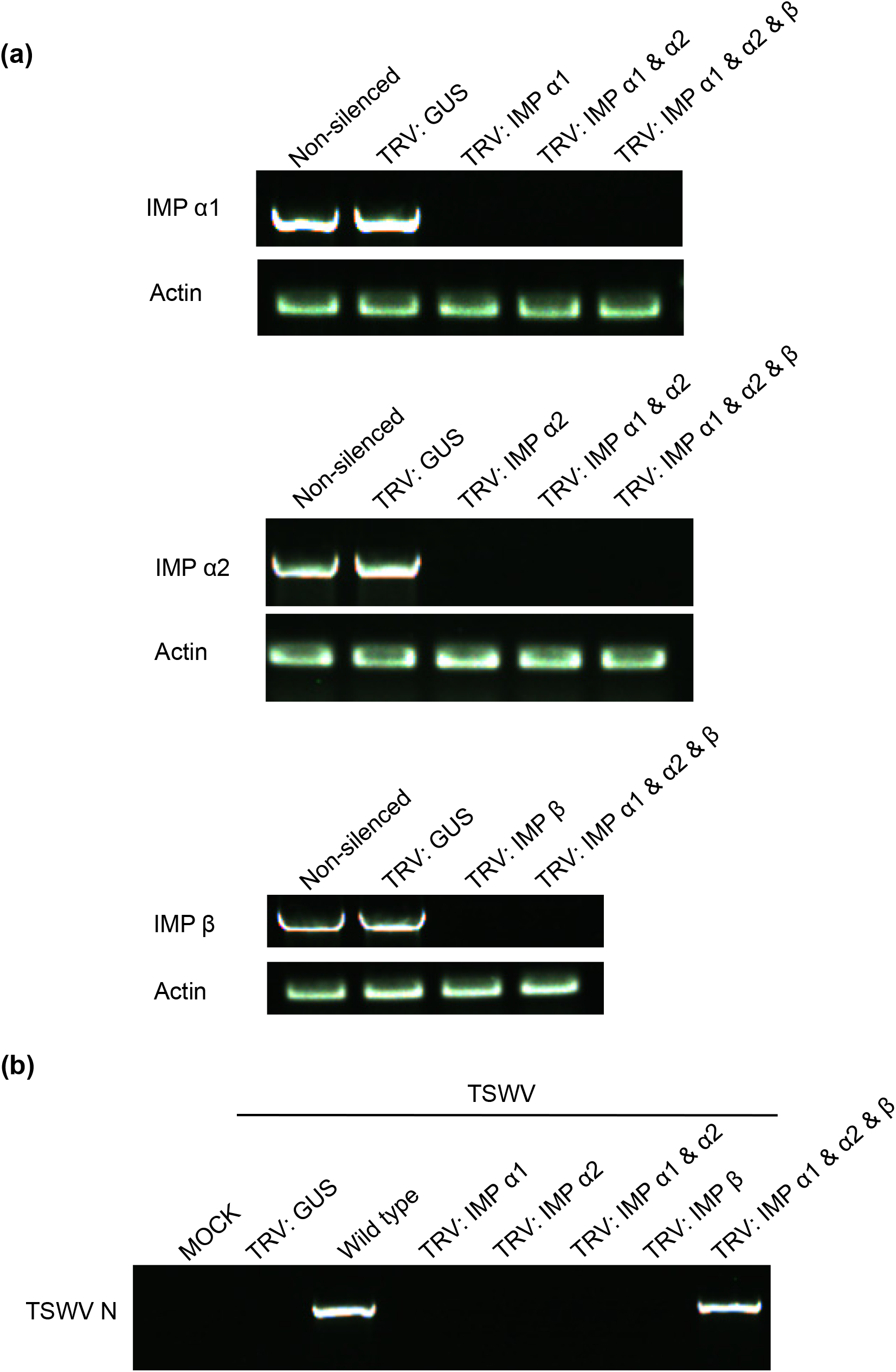
RT-PCR analyses of *importin* α*1*, α*2* and *β* expressions in the assayed plants and their effects on TSWV systemic infection. (a) Expressions of *importin* α*1,* α*2*, and *β* in Sw-5b transgenic *N. benthamiana* plants were silenced individually or together using a TRV-based VIGS vector. The gene silencing results were determined through semi-quantitative RT-PCR using gene specific primers. PCR products obtained after 25 cycles of PCR reaction were visualized in 1% agrose gel through electrophoresis. (b) RT-PCR detection of TSWV systemic infection in the assayed plants. The resulting PCR products were visualized in 1% agrose gel through electrophoresis.

**Table S1.** List of primers used in this study.

**Table S2.** Response of six different types of transgenic *Nicotiana benthamina* plants driven by 35S promoter to TSWV infection.

**Table S3.** Response of six different types of transgenic *Nicotiana benthamina* plants driven by Sw-5b native promoter to TSWV infection.

**Table S4.** Response of six different types of transgenic *Nicotiana benthamina* plants to TSWV infection.

**Table S5.** Mass spectrum data of YFP-SD

**Table S6.** Mass spectrum data of YFP-Sw-5b

